# Successful placentation in human pregnancy is regulated by reciprocal interactions between maternal uterine NK cells and fetal placental trophoblast

**DOI:** 10.1101/2023.06.07.544122

**Authors:** Qian Li, Andrew Sharkey, Megan Sheridan, Elisa Magistrati, Anna Arutyunyan, Oisin Huhn, Carmen Sancho-Serra, Holly Anderson, Naomi McGovern, Laura Esposito, Ridma Fernando, Lucy Gardner, Roser Vento-Tormo, Margherita Yayoi Turco, Ashley Moffett

**Author notes:** These authors contributed equally (Qian Li, Andrew Sharkey, Megan Sheridan, Elisa Magistrati). Correspondence (R.V.-T.), (M.Y.T.), (A.M.).

## Abstract

Fetal growth and development during human pregnancy depends on delivery of adequate maternal oxygen and nutrients to the fetus via the placenta. In humans, the balanced invasion of fetal placental trophoblast cells into the maternal uterine lining, where they interact with uterine natural killer cells (uNK), is thought to be critical for a successful pregnancy but exactly how this influences reproductive outcomes remains undefined. Here, we used our trophoblast organoid model and primary tissue samples to determine how uNK affect placentation. By locating potential interaction axes between primary trophoblast cells and uNK using single cell transcriptomics, and *in vitro* modelling of these interactions in trophoblast organoids, we identify a uNK-derived cytokine signal that promotes trophoblast differentiation by enhancing epithelial-mesenchymal transition and increasing trophoblast cells at the late stage of the invasive pathway. Moreover, it affects transcriptional programs involved in increasing blood flow, placental access to nutrients, and dampening inflammatory and adaptive immune responses, as well as gene signatures associated with disorders of pregnancy such as pre-eclampsia. Our findings shed new light on how optimal immunological interactions between maternal uNK cells and fetal trophoblast enhance reproductive success.

## Introduction

The placenta supports the fetus throughout pregnancy and abnormal placental development is a major cause of maternal and fetal mortality^1^. In humans, fetally-derived placental trophoblast cells develop from the trophectoderm surrounding the blastocyst and differentiate into two main lineages^2^. Cytotrophoblast cells cover the placental villi and form an overlying syncytium which mediates nutrient and gas exchange with maternal blood, while extravillous trophoblast cells (EVT) invade into the maternal decidual stroma, differentiating into interstitial EVT (iEVT) that eventually fuse to form placental bed giant cells (GC). iEVT migrate towards maternal spiral arteries and, in combination with endovascular EVT that moves down inside the arteries, remodel them to become high conductance vessels that can deliver a sufficient blood supply to the developing fetus^3^. Defects in this process can cause disordered blood flow into the intervillous space, damage to the placental villous tree and, thus, fetal growth restriction^4^. Abnormal placentation underlies common diseases of pregnancy including pre-eclampsia, stillbirth, fetal growth restriction and recurrent miscarriage^1,5^.

Even compared to other primates, the human placenta invades deeply into the decidual stroma; a likely reason for this is the requirement to ensure a sufficient blood supply necessary to support the *in utero* development of the large human brain^6,7^. This means that there are no suitable animal or *in vitro* models that adequately reflect the characteristic features of the human placenta. This has been a serious obstacle to understanding how EVT differentiation is regulated and why this process sometimes goes wrong. We have developed a trophoblast organoid system where cytotrophoblast cells can be differentiated to EVT^8^. Single cell RNA sequencing (scRNA-seq) and spatial transcriptomics of first trimester pregnant hysterectomy specimens shows that this model reliably recapitulates the *in vivo* EVT states^9^. As EVT invade the decidua they encounter maternal immune cells. There is a long standing notion that decidual leukocytes play a role in the pathogenesis of pre-eclampsia because there are immunological features of memory and specificity, demonstrated through epidemiological, genetic and functional studies^5^. The main population of immune cells that EVT will interact with are uterine Natural Killer (uNK) cells that are spatially and temporally associated with placentation^10–12^. Here, we use trophoblast organoids to investigate how uNK cells affect EVT differentiation and function.

Evidence that uNK affect placentation and the risk of developing pre-eclampsia has come from genetic studies of allogeneic interactions between Killer Immunoglobulin-like Receptors (KIR) on uNK and their HLA-C ligands expressed by EVT^13^. All HLA-C alleles can be assigned to C1 or C2 groups based on a dimorphism in the α1 domain. This means that EVT can express C1 and/or C2 epitopes. The highly polymorphic KIR family includes the *activating* KIR2DS1 and the *inhibitory* KIR2DL1 receptors that can recognise C2 epitopes on EVT^14,15^. Thus, each pregnancy is characterised by different combinations of maternal KIR and fetal HLA-C, resulting in variable inhibition or activation of uNK when they encounter EVT. Genetic studies suggest that mothers with KIR2DL1 but no KIR2DS1 and a C2+HLA-C fetus, are at increased risk of pre-eclampsia, while mothers with KIR2DS1 and fetal C2+HLA-C are protected^16–19^. Ligation of the *activating* KIR2DS1 stimulates uNK secretion of CSF2 (GM-CSF), which can influence trophoblast migration *in vitro*^17^ illustrating the functional relevance of uNK cytokines during placentation.

In this study, we use trophoblast organoids to investigate how uNK-derived cytokines influence EVT behaviour. By identifying cytokine-receptor pairs that mediate potential uNK-EVT interactions from *in vitro* modelling of KIR-HLA combination events and *in vivo* scRNA-seq profiling of maternal-fetal interface, we define a cocktail of uNK cytokines. Their functional consequences on EVT are investigated using trophoblast organoids followed by comparisons with primary tissue samples. This reductionist approach, by focussing on uNK-derived cytokines, overcomes the considerable intrinsic genetic and experimental difficulties of co-culturing first trimester primary uNK with EVT. Our findings show that cytokines derived from uNK act on EVT to regulate diverse processes important during early pregnancy. Combining trophoblast organoids with uNK cytokines provides an example of how this model system could be used to investigate how different decidual cell types regulate EVT functions required for successful placental development in humans.

## Results

### Identification of cytokine-receptor pairs that mediate interactions between uNK and EVT

NK cells in peripheral blood function by killing target cells or secreting cytokines/chemokines^20^. In contrast, uNK cells are poorly cytolytic but do produce cytokines that are likely to affect trophoblast behaviour^10^. Our previous studies using scRNA-seq and mass cytometry defined three uNK subsets, uNK1, uNK2 and uNK3^21,22^. Among these three uNK subsets, uNK1 and some uNK2 express different combinations of *activating* or *inhibitory* KIR that bind the C1+ and C2+HLA-C groups expressed by EVT^21^ (Figure 1A). These interactions affect cytokine production by uNK^17,23^. To test this further, we refined our previous experiment and modelled KIR-HLA interactions by co-culturing uNK (n=3 donors) with the HLA-null cell line, 721.221 (221), transfected to express either C1+ or C2+HLA-C allotypes. After stimulation using C1+ or C2+221 cells, uNK were separated using flow cytometry into three subsets: *activating* KIR2DS1 single positive (sp), *inhibitory* KIR2DL1sp, and KIR2DS1/KIR2DL1 double positive (dp) (Figure 1B). By transcriptional profiling of each subset using RNA-sequencing, KIR2DS1sp uNK (the protective activating KIR for C2+HLA-C), show distinct responses to C2+HLA-C compared to those seen with KIR2DL1sp and KIR2DS1/L1dp from the same donor (Figures S1A-S1C). Genes exclusively upregulated in KIR2DS1sp uNK are particularly enriched for cytokines such as *XCL1* and *CSF2* (Figures 1C and S1D). To validate whether these cytokines are released by all uNK subsets (uNK1, uNK2 and uNK3) *in vivo*, we examined the expression of common cytokines/chemokines in different cell types at the maternal-fetal interface using our previous scRNA-seq data^21^ (Figure S2A). This revealed four cytokines that are restricted to uNK cells in comparison with other decidual cell types: *XCL1*, *CSF2*, *CSF1*, and *CCL5* (Figure 1D). Among them, XCL1 and CSF2 are upregulated in KIR2DS1sp uNK after interaction with C2+HLA-C, confirmed by intracellular FACS (n=6 donors) (Figures 1E and S2B). Although CSF2 is only expressed at low levels by the uNK in the scRNA-seq data^21^ (Figure 1D), its detection at the protein level suggests it is possibly pre-synthesized and stored in granules (as occurs in eosinophils) to explain its rapid secretion after KIR activation^24^. CSF1 is a major product of uNK cells^21,25–27^ and CCL5 is made mainly by uNK3, demonstrated by both our scRNA-seq data and intracellular FACS (Figures 1D, 1F, and S2C). CSF1, CCL5 and XCL1 were predicted to interact with EVT in our previous scRNA-seq and spatial transcriptomic analysis of the maternal-fetal interface (CSF2 mRNA levels are low in *in vivo* scRNA-seq data)^9,21^. These interactions are further reinforced by the presence of the cognate receptors for all four cytokines in EVT, as demonstrated by immunohistochemistry of EVT *in vivo*^23,28–30^, our scRNA-seq data^21^ (Figure S2D), and by flow cytometry of primary trophoblast cells (Figure 1G). We have therefore identified four cytokines - CSF1, CSF2, XCL1, CCL5 - with restricted production by uNK whose receptors are expressed by EVT.

**Figure 1.**
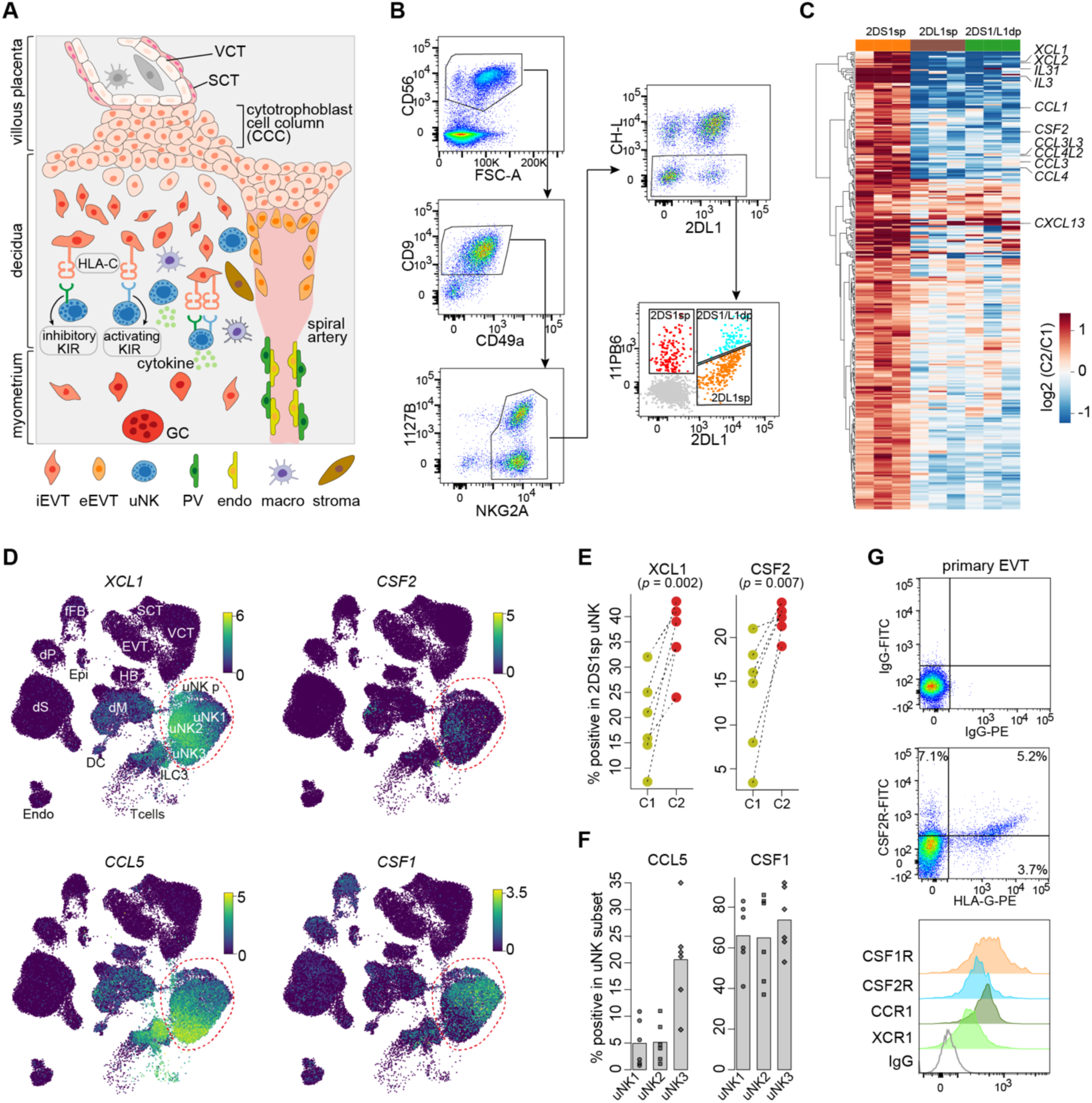
Identification of uNK-restricted cytokines that have receptors on EVT. (**A**) Diagram showing key cell types in the first trimester placenta, with interactions between maternal uNK and fetal EVT shown. VCT, villous cytotrophoblast; SCT, syncytiotrophoblast; GC, placental bed giant cells; iEVT, interstitial EVT; eEVT, endovascular EVT; PV, perivascular cells; endo, endothelial cells; macro, macrophages; p, proliferative. (**B**) FACS gating strategy for isolating the three uNK subsets: KIR2DS1 single positive (sp), KIR2DL1sp, KIR2DS1 and KIR2DL1 double positive (dp) after co-culture with 221-target cells expressing Cw*0501 (C2+HLA-C) or Cw*0802 (C1+HLA-C). (**C**) Heat map showing the log2-transformed fold change between culture with 221-C2+HLA-C and that with 221-C1+HLA-C targets across the three donors in each uNK subset for protein-coding genes specifically upregulated in the KIR2DS1sp subset (n=3 decidua; FDR < 0.05, and fold change > 1.5). (**D**) UMAP visualizations of the expression of the four uNK-restricted cytokines in different cell populations at the maternal-fetal interface based on our previous scRNA-seq data. dM, decidual macrophages; dS, decidual stromal cells; Endo, endothelial cells; Epi, epithelial glandular cells; dP, perivascular cells; DC, dendritic cells; fFB, fetal fibroblasts; HB, Hofbauer cells. (**E**) Intracellular staining of KIR2DS1sp uNK for cytokines XCL1 and CSF2 after 5 hours co-culture with 221-C2+HLA-C compared with 221-C1+HLA-C targets (percentage of positive staining cells in KIR2DS1sp subset compared to isotype-matched control, n=6). *p*-values were obtained from one-sided paired Student’s *t*-test. (**F**) Intracellular staining of CCL5 and CSF1 by flow cytometry in uNK1, uNK2 and uNK3 subsets after 5 hours culture with 221-C1+ target cells (similar results were obtained after co-culture with 221-C2+ targets (data not shown)). (**G**) Surface staining by flow cytometry for cytokine receptors CSF1R, CSF2Ra, CCR1 and XCR1 on freshly isolated EVT from first trimester decidual samples (representative staining of *n=3* donors). EVT were identified as HLA-G+ cells.

### Modelling uNK-EVT interactions using trophoblast organoids

To determine the effect of the uNK cytokine cocktail on EVT behaviour, we used our trophoblast organoid system^8^. We performed flow cytometry to confirm expression of the cognate receptors for these cytokines on organoids differentiated to EVT (Figure 2A), demonstrating the feasibility of modelling uNK-EVT interactions using organoids. To mimic the *in vivo* decidual microenvironment, we exposed the organoids to the uNK cytokine cocktail during the induction of trophoblast cells to EVT in EVT differentiation medium (EVTM) (Figure 2B). As a control, the uNK cytokines were also added to organoids cultured in trophoblast organoid medium (TOM) without any EVT differentiation (Figure 2B). We noticed that in some differentiating organoids cultured with these cytokines more cells appeared to be invading the Matrigel droplet from the organoid (Figure 2C). We therefore checked the expression of genes defining all different trophoblast sub-populations by reverse transcription polymerase chain reaction (RT-PCR) (Figure 2D). There was reduction in expression of genes characteristic of villous cytotrophoblast (VCT), *EPCAM*, *ITGA6* and *Ki67* (EVT no longer proliferate after beginning the invasive process), and upregulation of *HLA-G*, the definitive EVT marker, confirming differentiation to EVT^31–33^. ITGA2 is found in cells in a niche in the extravillous cytotrophoblast cell columns (CCC)^34^, and its expression is increased after addition of cytokines for 96h suggesting that they enhance EVT differentiation.

**Figure 2.**
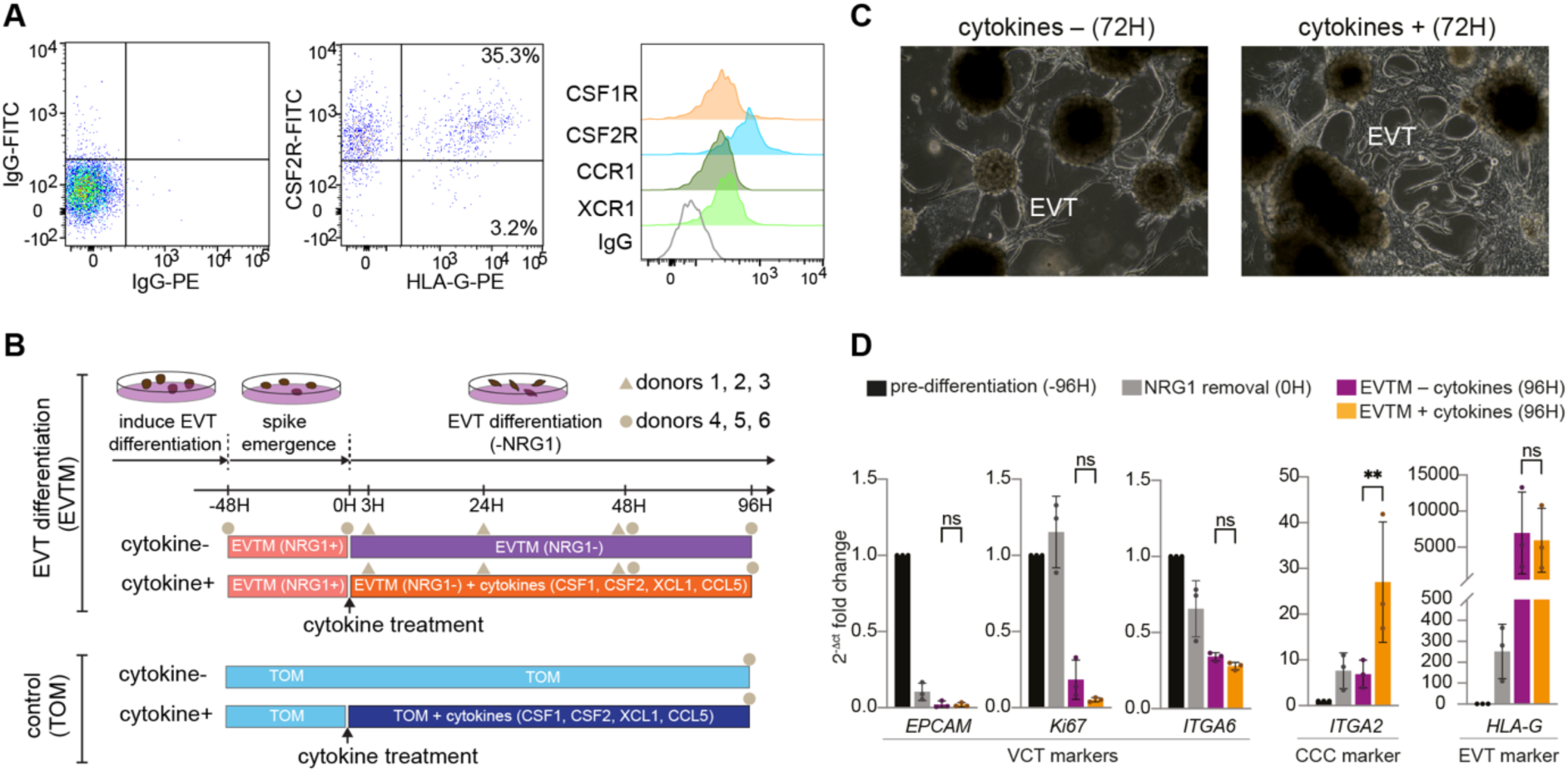
Modelling uNK-EVT interactions using trophoblast organoids. (**A**) Surface staining by flow cytometry for cytokine receptors CSF1R, CSF2Ra, CCR1 and XCR1 on EVT derived from trophoblast organoids differentiated to EVT (representative staining of *n=2* biological replicates). EVT were identified as HLA-G+ cells. (**B**) Experimental outline for trophoblast organoid differentiation and cytokine treatment, as well as single-cell transcriptome profiling at different time points from six donors. (**C**) Phase-contrast images of trophoblast organoids plated in Matrigel drop and differentiated to EVT with and without addition of a cocktail of selected cytokines: CSF1, CSF2, CCL5 and XCL1. (**D**) RT-PCR of trophoblast sub-population markers performed in trophoblast organoids at different time points during EVT differentiation, and treated with or without cytokines. Results are expressed as fold change of 2^-Δct^ with respect to the pre-differentiation condition. Bars represent mean ± SD (n=3 independent experiments). ns, not significant; \*\**p* < 0.005 by ratio paired *t*-test.

### Cytokines derived from uNK enhance EVT differentiation

To investigate further the cellular and molecular changes induced in EVT by the uNK cytokines, we performed scRNA-seq of organoids at different time points during EVT differentiation treated with and without cytokines (Figure 2B). After integration across organoids derived from different donors, followed by stringent quality control (Figures S3A-S3F), we obtained 67,996 cells which, based on the canonical marker genes, cover the three main trophoblast populations: VCT, syncytiotrophoblast (SCT) and EVT (Figure 3A). EVT were further subdivided into three early, two intermediate, and three late subtypes based on their gradually increasing expression of established EVT marker genes (Figure S3G). VCT and SCT are detected in both TOM and EVTM, while, as expected, EVT are only present in the latter (Figure 3A). To delineate the course of EVT differentiation, we next performed trajectory analysis by using the transcriptomic vector field in scTour^35^. This recapitulates the known bidirectional differentiation pathways from VCT towards either SCT or EVT, with EVT undergoing further continuous progression from early, intermediate to late stages (Figure 3A). At later time points there is a trend for an increased proportion of late EVT subtypes in cytokine-treated organoids (Figures 3B and S3H). For instance, EVT_late_3 is mainly detected at 96h from the organoids treated with cytokines in two donors (Figures 3B and S3F). This is reinforced by a differential abundance analysis which demonstrates that late EVT subtypes, especially EVT_late_3 are significantly enriched for cytokine-treated versus untreated cells (Figure 3C).

**Figure 3.**
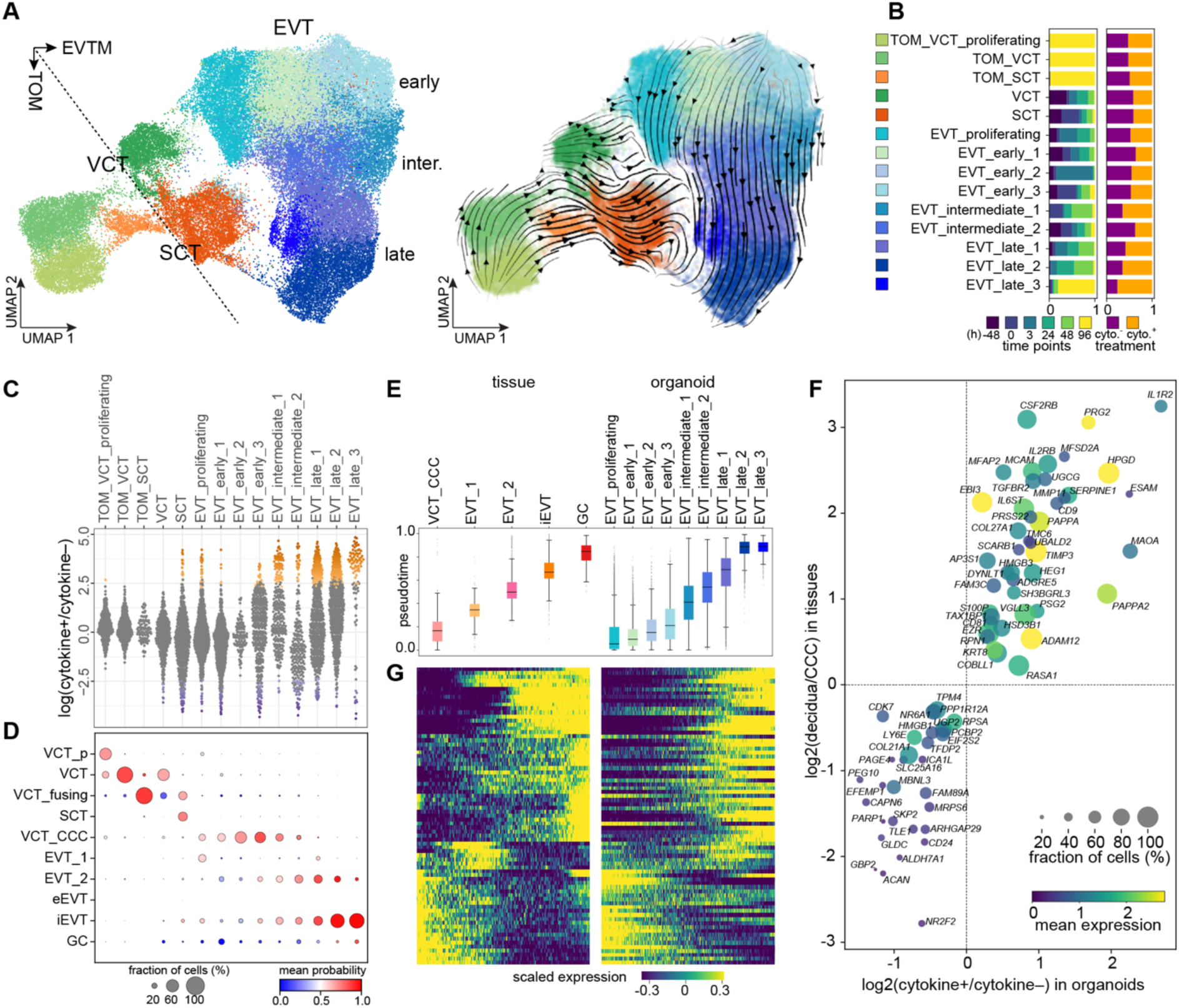
Trophoblast organoids treated with uNK cytokines mirror the cellular and molecular changes along the EVT differentiation pathway (A) UMAP visualisations of the 67,996 cells from trophoblast organoids with colours indicating cell types (left), and the streamplot showing the transcriptomic vector field (right). **(B)** The distribution of cells collected from different time points (left) and treatment conditions (right) in each cell type. (**C**) Cell abundance changes induced by the cytokine treatment. Dots represent neighbourhoods grouped by cell types, with coloured ones showing significant abundance changes (FDR < 0.1). (**D**) Predicted identities of the *in vitro* organoid cells based on the reference from the *in vivo* cells. Colour of dots indicates the prediction probability and dot size denotes the proportion of cells within a cell type assigned to the reference cell types. VCT_p, proliferative VCT; VCT_CCC, cytotrophoblast cell column VCT. (**E**) Box plot of the developmental pseudotime inferred for the *in vivo* cellular states and predicted for the *in vitro* ones based on the same scTour model. The medians, interquantile ranges, and 5th, 95th percentiles are indicated by centre lines, hinges, and whiskers, respectively. (**F**) Scatter plot showing genes consistently upregulated or downregulated after cytokine treatment in EVT_late_3 in organoids (x axis) and after invading the decidua (iEVT) compared to EVT located at the proximal end of the CCC (y axis). Colour shade and dot size denote the expression and proportion averaged between the *in vitro* and *in vivo* EVT, respectively. (**G**) The expression dynamics along the developmental pseudotime estimated in (**E**) for genes from (**F**).

Given the diverse cell subtypes detected in the organoids both treated with and without cytokines, we next asked how they mirror the cellular states of trophoblast *in vivo*. To resolve this, we aligned the *in vitro* cell types with primary trophoblast analysed from human implantation sites by single nucleus RNA sequencing (snRNA-seq)^9^. This reveals an *in vitro-in vivo* correspondence for all three trophoblast populations (Figure 3D); thus, VCT and SCT from organoids match their *in vivo* counterparts. For the organoid EVT subtypes, early ones mainly correspond to EVT progenitors (VCT_CCC)^9^, while intermediate ones span multiple *in vivo* states from these EVT progenitors, EVT located at the distal end of the cell column (EVT_2)^9^, to iEVT deeper in decidua (Figure 3D). Late EVT cells are largely assigned to iEVT, particularly EVT_late_3 which also includes a few cells predicted to be GCs and expressing GC marker genes (Figures 3D and S3G). These GCs form deep within the decidua and are the end-point of the EVT differentiation pathway^36^. As before, no endovascular EVT is observed in the organoids even after treatment with cytokines^9^ (Figure 3D). These *in vitro-in vivo* comparisons are reproducible when aligned to a different *in vivo* scRNA-seq dataset of the maternal-fetal interface^21^ (Figure S4) and are consistent with our recent benchmark of primary trophoblast organoids without cytokines^9^. To see how differentiation of EVT is affected by exposure to uNK cytokines, we performed unsupervised pseudotime ordering of both *in vivo* and *in vitro* EVT. The same trajectory is seen; EVT progenitors gradually differentiate to GCs in the primary tissue as previously shown^9^, whilst early EVT progressively differentiate to late EVT in organoids (Figures 3E and S5A). To summarise, the organoids cultured with uNK-derived cytokines recapitulate the EVT differentiation pathway seen *in vivo,* resulting in an increased proportion of late EVT, especially EVT_late_3 similar to iEVT *in vivo* (Figures 3C-3E).

### uNK-derived cytokines affect EVT behaviour by regulating diverse gene programs

uNK are present throughout the decidua and mainly interact with iEVT after they have left the CCC and infiltrated the uterus (Figure 1A). Because EVT_late_3 is the population that is particularly enriched after cytokine treatment, we focussed on this subtype to identify molecular changes occurring in EVT after exposure to cytokines. We used both our *in vivo* snRNA-seq^9^ (Figure 3F) and scRNA-seq^21^ (Figure S5B) datasets to also compare changes in the organoids with those seen between *in vivo* EVT located at the proximal end of the CCC (less accessible to uNK cytokines) and iEVT (fully accessible to uNK cytokines). We find 44 upregulated and 31 downregulated genes in both organoids after cytokine exposure and in primary trophoblast (Figures 3F, S5B and S5C) with consistent *in vitro*-*in vivo* temporal profiles along the EVT differentiation pathway (Figures 3G and S5A). Genes downregulated by cytokines are enriched in cell cycle control, consistent with the non-proliferative nature of iEVT (Figures 4A and S5D). Genes upregulated in EVT after exposure to uNK cytokines are involved in several different biological processes (Figure S5D). In line with the morphological and cellular changes supporting enhancement of EVT differentiation, genes involved in EMT (*MCAM*, *TGFBR2*, *HEG1*, *VGLL3*)^37–40^, cell invasion (*ADGRE5*, *MMP11*, *TIMP3*, *S100P*)^41–43^, and fusion (*CD9*, *ADAM12*)^44–46^ are up-regulated by uNK cytokines (Figures 4A and S5D). Additionally, a group of subunits of cytokine receptors (*IL2RB*, *CSF2RB*, *IL6ST*, *EBI3*, *IL1R2*) with their cognate ligands mainly expressed by decidual macrophages and uNK cells are positively regulated by cytokines (Figures 4A, S5D and S5E). Amongst them, IL1R2 acts as a decoy receptor for IL1B produced in abundance by decidual macrophages (Figure S5E), suggesting a mechanism for dampening local inflammation in the placental bed^47^. We also detect genes (*MAOA*, *PRG2*, *PAPPA*, *PAPPA2*) involved in placental access to nutrients and regulation of blood flow. PAPPA and PAPPA2, together with ADAM12, cleave insulin-like growth factor binding proteins (IGFBP), thus regulating the availability of IGFs^48–50^. To test this, we collected organoid supernatants under different treatment conditions and confirm that IGFBP levels are indeed reduced with cytokines (Figure S5F). MAOA encodes an enzyme that degrades serotonin, which is vasoactive^51^, and once again we find decreased levels of serotonin in cytokine-treated organoid supernatants (Figure S5G).

**Figure 4.**
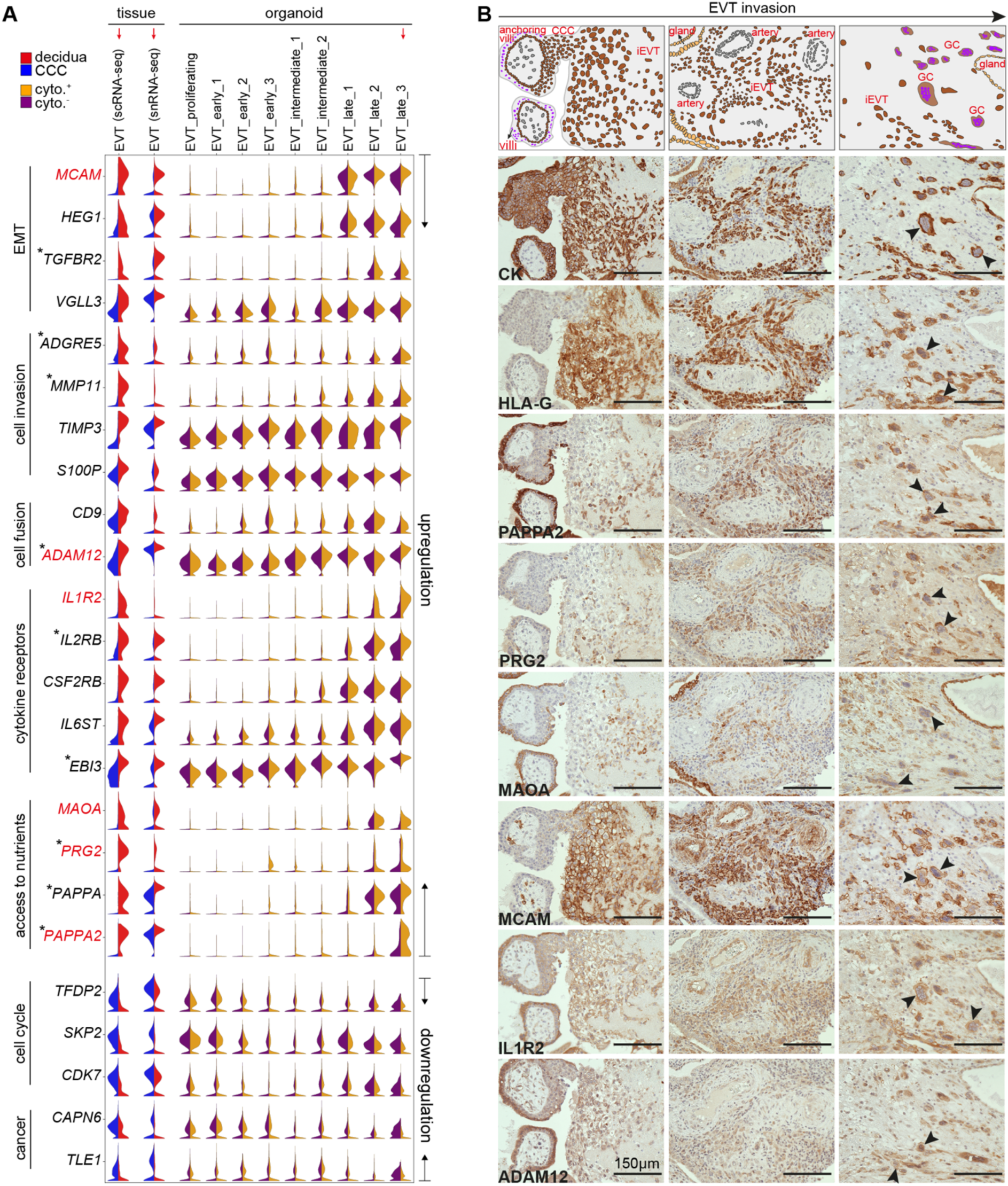
uNK cytokine-induced molecular changes in EVT (A) Selected molecular changes induced by the uNK cytokines, with the expression patterns in *in vivo* EVT cells collected from decidua (red) and CCC (blue) shown in the first two columns, and in different EVT subtypes from organoids treated with (orange) and without (purple) cytokines in the following columns. Genes marked by asterisks are associated with a risk of pre-eclampsia or other reproductive disorders. (**B**) Immunohistochemistry of serial sections of an anchoring villus (left), iEVT encircling a spiral artery (middle) and placental bed giant cells (right) for proteins highlighted in red in (**A**) together with Cytokeratin (CK) and HLA-G. Arrows indicate giant cells. Top panel shows decidua basalis at different depths indicated by the Cytokeratin staining.

To verify expression of these genes along the EVT differentiation pathway *in vivo*, we performed immunohistochemistry on serial sections of first trimester implantation sites (Figures 4A and 4B). The sections spanned the entire decidua basalis from the anchoring villi, iEVT encircling spiral arteries, to placental bed GCs present close to the decidual-myometrial border (Figure 4B). PAPPA2, PRG2, MAOA, MCAM, and IL1R2 are absent or only weakly expressed in CCC of the anchoring villi but are expressed by iEVT (Figure 4B), consistent with their profiles from scRNA-seq (Figure 4A). MCAM expression is striking with no staining in CCC but strong positive staining seen at the distal end of the cytotrophoblast shell at the interface with decidua, whereas expression of PAPPA2, PRG2, and MAOA first appears on iEVT deeper within the decidua (Figures 4A and 4B). GCs, the end point of EVT differentiation, stain positively for all proteins tested (Figure 4B), providing further evidence that GCs are present in the organoids after exposure to uNK cytokines. ADAM12, IL1R2, PAPPA2 and MCAM are also expressed by endovascular trophoblast with scattered PRG2 and MAOA positive cells in the trophoblast plugs (Figure S6).

To further confirm that the changes observed are indeed induced by uNK-derived cytokines, we performed RT-PCR in organoids treated with or without cytokines to examine expression differences for genes representative of different biological processes. In keeping with our scRNA-seq data, expression of all genes analysed is induced in the first phases of EVT differentiation, and further increased (for upregulated genes) or decreased (for downregulated genes) with cytokine treatment (Figure 5A). We next sought to verify if these changes occurred in invading EVT migrating out of the organoid structure. To do this, we specifically isolated only migrating EVT and obtained an *HLA-G*-enriched population that lacks the VCT marker, *EPCAM*, and the SCT marker, *ERVW1* (Figure S7A). Quantification by RT-PCR of these purified EVT cells confirms the enrichment of the transcriptional changes induced by uNK cytokines in more differentiated EVT (Figure 5B). To validate these findings at the protein level, we used immunofluorescence to quantify two representative upregulated genes: MAOA and MCAM. Both show increased levels of immunostaining in migrating EVT (Figure S7B) in the presence of cytokines (Figure 5C).

**Figure 5.**
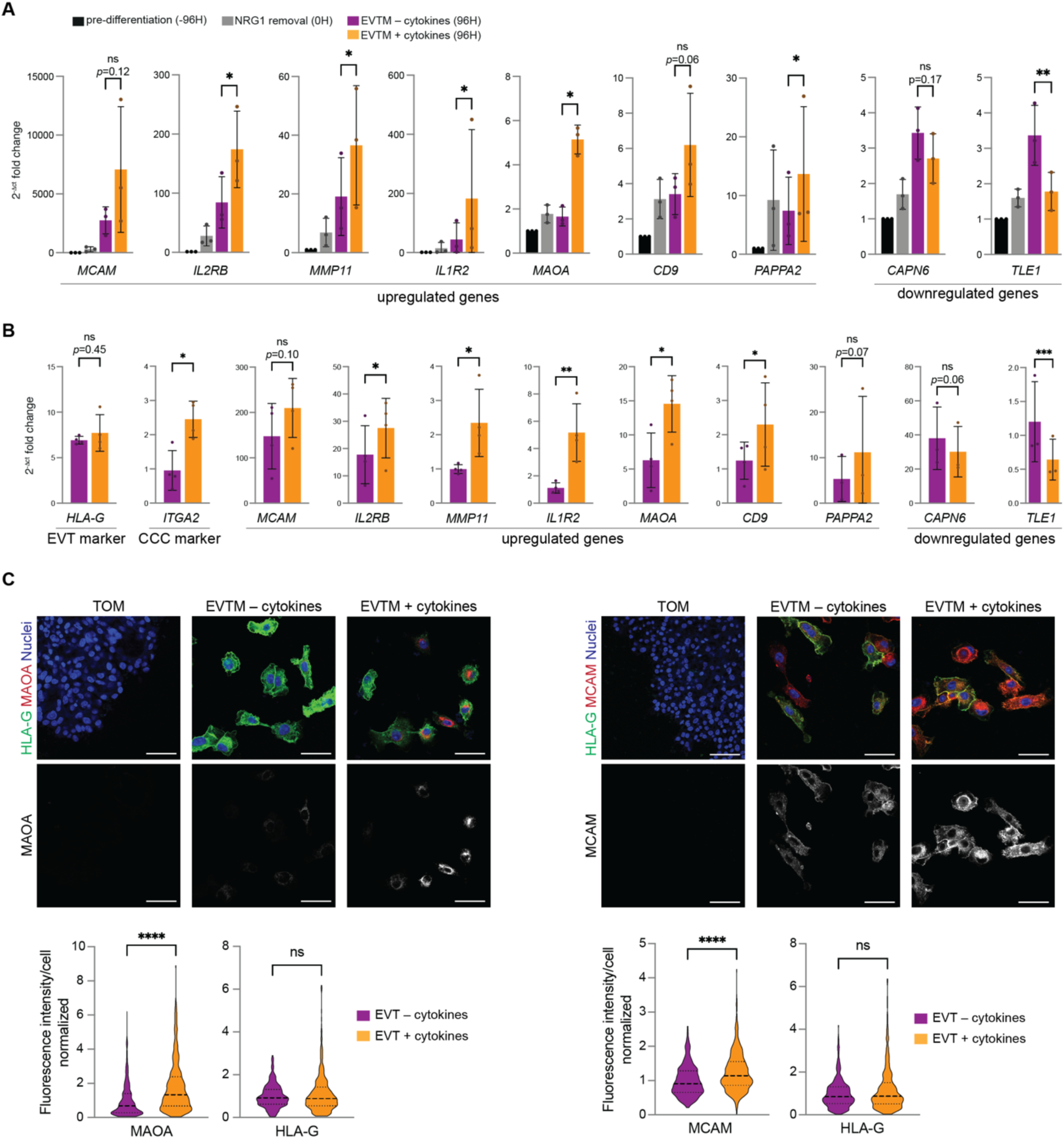
Verification of molecular changes after uNK-derived cytokine treatment in organoids. (A) RT-PCR of selected genes found as up- or down-regulated in cytokine-treated organoids upon scRNA-seq. The RT-PCR was performed in trophoblast organoids at different timepoints during EVT differentiation, either treated or not treated with cytokines. Results are expressed as fold change of 2^-Δct^ with respect to the pre-differentiation condition. Bars represent mean ± SD (*n*=3 independent experiments). *ns*, not significant, \**p*<0.05, \*\**p*<0.005 by ratio paired *t*-test. (**B**) RT-PCR of genes selected as in (**A**) and trophoblast markers *HLA-G* and *ITGA2*. The RT-PCR was performed in EVT cells isolated from organoids after 96h from NRG1 removal, treated with and without cytokines. Results are expressed as 2^-Δct^. Bars represent mean ± SD (*n*=3 or 4 independent experiments). *ns*, not significant, \**p*<0.05, \*\**p*<0.005, \*\*\**p*<0.0005 by ratio paired *t*-test. (**C**) IF analysis of MAOA and MCAM protein expression in organoids treated with or without cytokines. Organoids were fixed and immunostained with the indicated antibodies after 96h from NRG1 removal. Top: representative images of three independent experiments, scale bars 50 μm. Bottom: Quantification of MAOA, MCAM and HLA-G intensity in isolated cells. Results are expressed as fold change with respect to non-treated average intensity and represented with violin plot reporting median and quartiles. MAOA and HLA-G (left): -cytokines, *n*=371 cells; +cytokines, *n*=410 cells from three independent experiments. MCAM and HLA-G (right): -cytokines, *n*=405 cells; +cytokines, *n*=378 cells from three independent experiments. *ns*, not significant, \*\*\*\**p*<0.0001 by Mann-Whitney test.

Our findings reveal the importance of the pathways we detect in EVT (Figure 6). Indeed, many of the genes we find upregulated in EVT by uNK cytokines are associated with increased risk of pre-eclampsia or other reproductive disorders (Figure 4A; Table S1).

**Figure 6.**
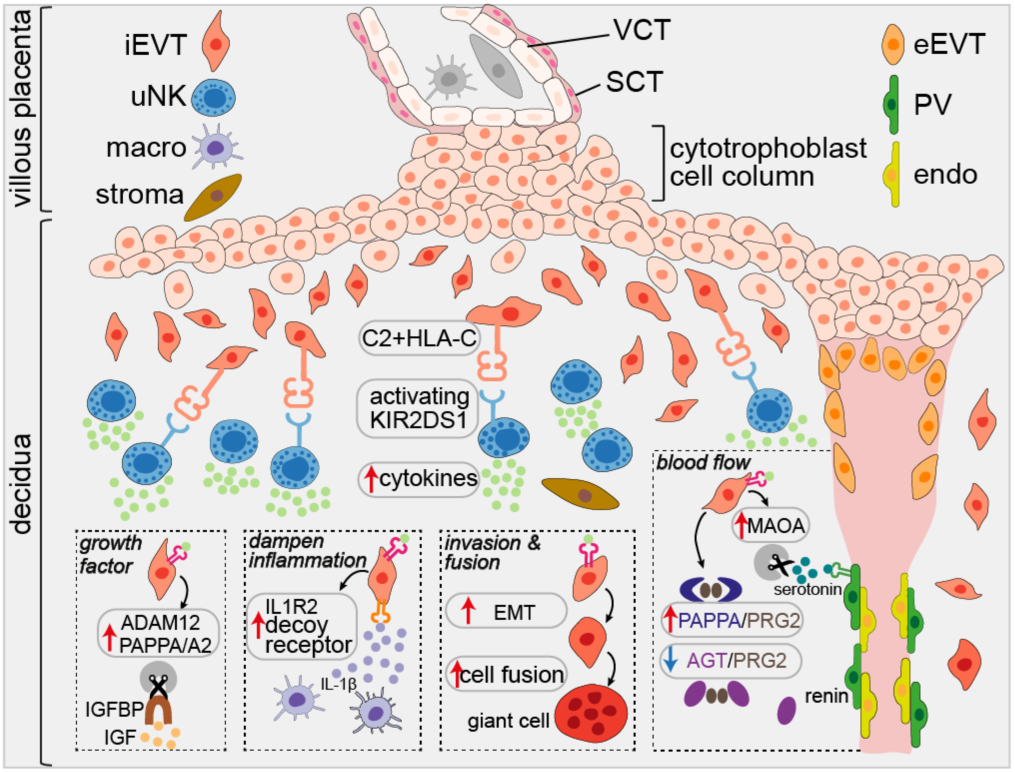
Schematic representation of the reciprocal interactions between uNK cells and EVT. Binding of the uNK cytokines to EVT triggers multiple effects including availability of growth factors, dampening of inflammation, enhancing EVT invasion and fusion, and influencing blood flow. VCT, villous cytotrophoblast; SCT, syncytiotrophoblast; iEVT, interstitial EVT; eEVT, endovascular EVT; uNK, uterine natural killer cells; macro, macrophages; PV, perivascular cells; endo, endothelial cells; EMT, epithelial to mesenchymal transition.

## Discussion

The primary pathogenesis of major disorders of pregnancy, including pre-eclampsia, is defective placentation and arterial transformation by EVT^52^. Because of the inaccessibility of the maternal-fetal interface early in gestation, and the ethical and logistical issues in obtaining samples, the underlying mechanisms responsible remain largely unknown. Here, we have taken advantage of trophoblast organoids differentiated to EVT that we have shown recapitulate the diverse cellular states of EVT *in vivo*^9^. These organoids provide a new model to investigate how interactions between maternal decidual cells and fetal placental EVT early in pregnancy influence reproductive outcomes. Although uNK cells amass at the site of placentation early in human pregnancy, it is not yet possible to mimic this *in vitro* because of the technical limitations involved in co-culturing primary uNK with trophoblast organoids. Therefore, the reductionist approach we have taken here is to focus on cytokine-receptor pairs that mediate uNK-EVT interactions, with the cytokines (XCL1, CSF2, CSF1, and CCL5) predominantly and specifically produced by uNK and the receptors expressed by EVT. Although, based on our scRNA-seq data, *XCL1* and *CCL5* are also expressed by some T cells, these are present in small numbers during the first trimester, and are sparse at the site of placentation^53^. Thus uNK will be the primary source of these cytokines.

We exposed trophoblast organoids differentiating to EVT to this cocktail of uNK cytokines, and analysed their effects on the developing placenta by comparing single cell transcriptomic data profiling trophoblast differentiation in both the organoids and primary tissue. We find that uNK cytokines enhance EVT differentiation by increasing the abundance of more differentiated EVT cells and regulating gene pathways involved in epithelial-mesenchymal transition, cell invasion and fusion. In our previous benchmarking of the trophoblast organoids, no GC were found^9^. In contrast, with the addition of uNK cytokines, both the transcriptomic alignment between cell types *in vivo* and *in vitro*, and the expression of GC signature genes validated in organoids and spatially in primary tissue, indicate the presence of terminally differentiated GC. Thus, with the uNK cytokine cocktail, the organoids recapitulate the full progenitor EVT-iEVT-GC *in vivo* differentiation pathway. Although it has long been suspected that uNK affect EVT invasion, results from previous studies have been conflicting^54–63^. This is probably because, in the invasion assays used, the identity and purity of the trophoblast cells is questionable. In addition, information on maternal KIR and fetal HLA-C genotypes is lacking. Our findings with 3D trophoblast organoids, in combination with transcriptome profiling at single-cell resolution, provide solid evidence supporting the positive role of uNK cytokines in EVT differentiation.

Apart from promoting differentiation, other interesting and unexpected roles for uNK cytokines in affecting EVT behaviour emerge from the transcriptional programs altered after cytokine exposure in the organoids. These signatures parallel those seen along the EVT differentiation pathway *in vivo,* validated in the primary tissue samples and organoids at the mRNA and protein levels, suggesting their comparable roles *in vitro* and *in vivo.* For instance, a group of subunits of cytokine/chemokine receptors are enriched after exposure to cytokines. Among them, IL1R2 binds IL1B (produced by decidual macrophages) but does not signal^64–66^, ensuring that all inflammatory IL1B would be mopped up at the site of placentation. Another cytokine receptor, IL2RB, may exist as a soluble form that could interact with any IL2 present to avoid T cell activation in decidua^67^. This could also be prevented by the upregulation of *VGLL3* which acts as a transcriptional regulator to drive PDL1/2 expression^68^, one potential mechanism for decidual T cell tolerance. Thus, these findings reinforce our earlier results that uNK have an essential role in dampening any innate or adaptive immune responses during placentation^21^. Another potential role for uNK cytokines is to increase blood flow and access to nutrients to ensure optimal fetal growth and development. We propose a mechanism whereby uNK cytokines increase the expression of ADAM12, PAPPA, and PAPPA2 that act to cleave IGFBPs to increase the availability of insulin-like growth factors^48–50^. Delivery of nutrients is also likely to be affected, indicated by increased expression of MAOA, which degrades EVT-derived serotonin and could therefore alter uterine blood flow^51^. Further evidence for this comes from the upregulated complex formed by PRG2 and PAPPA/PAPPA2 which competes with PRG2/angiotensinogen complexes^69^, thus releasing angiotensinogen to participate in the renin-angiotensin system for delivery of maternal blood to the placenta. Further work could focus on the detailed underlying mechanisms of these pathways in human pregnancy, and dissecting out effects of individual cytokines.

Previous efforts to understand the pathogenesis of disorders of pregnancy have resulted in identification of a variety of candidate genes and serum biomarkers. Our study does also provide a link between the effect of uNK cytokines on placentation with disorders including pre-eclampsia, fetal growth restriction and placenta accreta syndrome. Firstly, secretion of two uNK cytokines (CSF2 and XCL1) is increased after the activating KIR2DS1 binds to C2+HLA-C ligand; this genetic combination of mothers with KIR2DS1 and fetal C2+HLA-C is protective against pre-eclampsia and associated with larger babies^17,18,70^. Secondly, several of the gene signatures we have found altered by uNK cytokines have already been associated with these disorders in several studies: *EBI3*, *MMP11*, *IL2RB*, *TGFBR2*, *ADAM12*, *PRG2*, *PAPPA* and *PAPPA2*^71–86^ (Table S1). Our study highlights the importance of now investigating the underlying mechanisms regulated by these genes that lead to different reproductive outcomes in humans, and provides candidate pathways with potential translational links to the clinic.

## Acknowledgements

The authors are grateful to patients for donating tissue for research. We thank staff at the Addenbrooke’s Hospital, Cambridge and at the West Suffolk Hospital, Bury St Edmunds and J. Bauer, A. Karcanias, P. Howden and C. Reitter from Cambridge Genomic Services, Department of Pathology. Special thanks to the Flow Cytometry Core Facility at the Department of Pathology and Core staff at the Immunophenotyping Hub at the Department of Medicine (University of Cambridge). 721.221 cells expressing either HLA-C*05:01, or HLA- C*08:02 were the kind gift of Malcolm Sim. A. Moffett holds a Wellcome Trust investigator award (200841/Z/16/Z). M.Y. Turco was supported by the Royal Society Dorothy Hodgkin Fellowship (DH160216) and Royal Society Research Grant (RGF\R1\180028) and has received funding from the European Research Council (ERC) under the European Union’s Horizon 2020 research and innovation programme (Grant agreement no. [853546]). N. McGovern is funded by a Wellcome Trust Sir Henry Dale and Royal Society Fellowship (grant 204464/Z/16/Z). A. Sharkey is funded by the Medical Research Council (grant MR/ P001092/1). R. Vento-Tormo was supported by Wellcome Sanger core funding (WT206194).

## Author contributions

M.Y.T. and A.M. conceived and designed the study. A.S. performed the uNK/HLA-C experiments and analysed the data with help from N.M., O.H. and L.E. Trophoblast organoid culture was performed by E.M., H.A., L.G., M.S. M.S. and R.F. performed cell collections from organoids for scRNA-seq. C.S.S performed scRNA-seq experiment. Q.L. analysed and interpreted the data. A.A. assisted with data analysis. E.M. performed RT-PCR and IF experiments in organoids. Q.L. performed IHC on tissue sections. R.V.-T., M.Y.T. and A.M. assisted in the interpretation of results. Q.L., A.S., E.M., M.Y.T and A.M. wrote the manuscript.

## Declaration of interests

The authors declare no competing interests.

## Methods

### RESOURCE AVAILABILITY

#### Lead contact

Further information and requests for resources and reagents should be directed to and will be fulfilled by the lead contact, Ashley Moffett.

#### Materials availability

Materials generated in this study are available upon reasonable request.

#### Data and code availability

All the data generated in this study are being uploaded to EMBL-EBI ArrayExpress. Codes used for all the analyses will be available in https://github.com/LiQian-XC.

### EXPERIMENTAL MODEL AND SUBJECT DETAILS

#### Human samples

Matched peripheral blood, decidual or placental samples were obtained from healthy women undergoing elective terminations in the first trimester of pregnancy (total numbers of donors, n=19, Table S3). Written and informed consent was obtained in accordance with the guidelines in The Declaration of Helsinki 2000. Ethical approval for the use of these tissues was obtained from the Cambridge Local Research Ethics Committee (REC 04/Q0108/23). This now forms part of the Centre for Trophoblast Research biobank for the ‘Biology of the Human Uterus in Pregnancy and Disease, Tissue Bank’ at the University of Cambridge. Overall biobank ethical approval is from the East of England–Cambridge Central Research Ethics Committee (17/EE/0151).

### METHOD DETAILS

#### Cell culture

Decidual mononuclear cells were routinely isolated by enzymatic digestion of decidual tissue as described previously, followed by density gradient centrifugation through Pancoll (PAN-Biotech) and either used fresh, or cryopreserved in 10%DMSO/90%FCS^22^.

Trophoblast organoids were obtained by sequential digestion of placental villi in 0.2% trypsin (PAN-Biotech P10-025100P), 0.02% EDTA (Sigma E9884) in PBS, followed by collagenase (1.0 mg ml−1 collagenase V, Sigma C9263) in Hams F12/10% FBS. Pooled digests were washed in Advanced DMEM/F12 medium, then resuspended in Matrigel on ice (Corning, 356231) and plated in 25 μl droplets into 48 well culture plates and overlaid with 250 μl trophoblast organoid medium^8^ (Table S2). Cultures were maintained in 5% CO2 in a humidified incubator at 37 °C and medium was replaced every 2–3 days. A list of the lines used in this work is available in Table S3.

#### Stimulation of uNK cells by HLA-C

uNK cells were stimulated by incubation with 721.221 cells expressing either HLA-C*05:01, which is a C2 allotype (C2+HLA-C) or HLA-C*08:02 which is a C1 allotype (C1+HLA-C). HLA-C*08:02 differs from HLA-C*05:01 by only the C1 and C2 epitopes (amino acids at p77 and p80) as previously described^87^. All 221 cells were cultured in RPMI 1640 medium (Gibco) with antibiotics, and 10% fetal calf serum (FCS).

#### Phenotyping to identify donors with uNK expressing KIR2DS1

For experiments to sort uNK for expression profiling, uterine NK cells from 4 donors were first phenotyped prior to co-culture, to determine their KIR2DL1 and KIR2DS1 expression. Freshly isolated decidual cells were isolated by mechanical sieving using a 70 micron sieve as described previously, since this provides better preservation of enzyme-sensitive epitopes required for immediate flow cytometry^88^. An aliquot of the resulting suspensions in PBS were immediately stained for viability with Zombie Aqua diluted 1:1000 (BioLegend) for 20 min at 4°C. Cells were washed twice in FACS buffer (PBS, with 2% FCS and 2 mM EDTA) and incubated with 10μl human AB serum in a final volume of 100μl of FACS buffer for 15 min to block nonspecific binding sites. Staining with a uNK phenotyping antibody panel was performed for a total of ∼35 min at 4°C. Antibodies used are listed in Table S4. To stain for KIR2DL1 or KIR2DS1 in the same sample, cells were stained with 3μl KIR2DL1 APC antibody (clone REA284; Miltenyi Biotec) in the cocktail with the other antibodies for 23 minutes. Then 7μl KIR2DL1/S1 (clone 11PB6; Beckman Coulter) was added for the final 12 minutes. After washing in FACS buffer, cells were fixed in 2% paraformaldehyde (Alfa Aesar, J61899). Analytical flow cytometry was performed using a Cytek Aurora spectral analyser (Cytek), and FACS data were analysed using FlowJo (Tree Star). 3 of the 4 tested donors were found to express KIR2DS1 and subsequently used for sorting of uNK.

#### Sorting of uNK to profile responses of KIR2DS1+ uNK after co-culture with C2+HLA-C

To determine responses of KIR2DS1+ uNK to C2+HLA-C, samples of DL from KIR2DS1+ donors (n=3) were co-cultured with 221 target cells expressing a C2+HLA-C allotype or C1+HLA-C as a control. After initial mechanical isolation and phenotyping as described above, decidual cell suspensions (10 million per flask), were cultured overnight to recover in 20 ml of RPMI 1640 with 10% heat-inactivated FCS and antibiotics, supplemented with 2.5 ng/ml IL- 15 (Peprotech). This dose of IL-15 maintains uNK viability without activating the cells. After 12 hours, non-adherent cells (largely DL) were recovered by vigorous washing in complete medium. The primary DL were stimulated by co-culture with 721.221 cells expressing either HLA-C*05:01 or HLA-C*08:02 at an effector target ratio of 1:1. Primary DL and target cells were co-cultured (each at 1.2×10^5^ per well) in a total volume of 700ul a 48 well flat bottomed plate in complete medium with 2.5 ng/ml IL-15. Plates were centrifuged for 2 min at 300 rpm followed by incubation 12hr at 37 °C, 5% CO2. Cells were recovered into FACS tubes (Falcon, 352054), washed in PBS and immediately viability stained with Zombie Aqua diluted 1:1000 (BioLegend) for 20 min at 4°C. Cells were then washed twice in FACS buffer, blocked and stained with the uNK cell sorting antibody panel as described above. After staining, subsets of CD56+CD3-CD9+CD49+ uNK expressing KIR2DS1 or KIR2DL1 were immediately purified by cell sorting into complete culture medium using a Becton Dickinson FACS Aria III controlled by BD FACS DIVA software (version 8). Details of antibodies used for cell sorting are listed in Table S5 and the gating strategy is shown in Figure 1B. Sorted cells were pelleted by centrifugation and frozen for subsequent RNA isolation using a total RNA isolation kit (Norgen, 51800).

#### Transcriptional profiling of KIR2DS1+ uNK responses to C2+HLA-C

RNA sequencing of sorted uNK subsets was performed at Cambridge Genomic Services, University of Cambridge using SMART-Seq v4 to generate the cDNA (Takara) and then adapter and index ligation was performed using the Nextera XT workflow (Illumina). Both were performed following manufacturer protocols. The resulting libraries were sequenced using a NextSeq 500 high output 75 cycles kit.

#### Intracellular staining for cytokine responses in uNK subsets

To verify cytokine responses in selected uNK subsets, frozen DL from KIR2DS1+ donors (n=6) were first thawed and recovered for 12hr in complete medium with IL-15. Non adherent cells were recovered. These primary DL were stimulated by co-culture with 721.221 cells expressing either HLA-C*05:01 or HLA-C*08:02 at an effector target ratio of 1:1 in 48 well plates as described above for sorting experiments. Plates were centrifuged for 2 min at 300 rpm followed by incubation 1hr at 37 °C, 5% CO2. GolgiPlug and GolgiStop (BD Biosciences, 555029 and 554724) were then added each at 1:500 dilution. After an additional 4 h of co-incubation at 37 °C, cells were recovered into FACS tubes for staining. To permit analysis of cytokine expression in both KIR2DS1/2DL1+ uNK as well other uNK subsets, an expanded antibody panel was employed (Table S6). The XCL1 antibody was directly labelled with CF594 using the Mix-n-Stain™Antibody Labelling Kit, (Biotium, 92256). After surface marker staining as described for sorts above, cells were fixed and permeabilized with BD Cytofix/Cytoperm™ kit (BD Biosciences, 554714) according to manufacturer’s instructions and stained with antibodies to the cytokines CSF2, XCL1, CSF1 and CCL5. Flow cytometry of the expanded 15 colour panel was performed using a Cytek Aurora.

#### Generation and analysis of EVT cells from trophoblast organoids by flow cytometry

The protocol for passaging and differentiation of trophoblast organoids has been described in detail by Sheridan et al. 2020^89^. In brief, after passaging into 35mm dishes, organoids are grown in TOM for 3-4 days then switched into EVT differentiation medium with NRG1 (EVTM; Table S7). After ∼7 days, when outgrowths of cells are observed from the organoids, the medium is switched to EVTM without NRG1 (EVTM-NRG1) for a further 7-10 days to complete EVT differentiation. To observe the effects of cytokines on EVT differentiation, cytokines are added to the EVTM-NRG1 medium as follows: CSF2 (10 ng/mL), XCL1 (100 ng/mL), CSF1 (20 ng/mL), and CCL5 (50 ng/mL). To analyse expression of HLA-G or cell surface receptors, by flow cytometry on the resulting EVT, the organoids are retrieved from Matrigel with Cell Recovery Solution (Corning 354253). After dissociation with 0.2% trypsin 250 (Pan Biotech P10-025100P), 0.02% EDTA (Sigma E9884) in PBS at 37 °C for 5 min, cells were washed in medium containing FBS and passed through a 40-μm cell strainer (Falcon 2340). Cells were blocked with human IgG (Sigma I4506) in Dulbecco’s PBS (ThermoFisher Scientific 14190136) with 1% FBS before labelling with W6/32–Alexa-488 anti-HLA-A, B, C antibody, HLA-G–PE, or isotype-matched controls (Table S8, antibodies for HLA-G and cytokine receptor staining). LIVE/DEAD Fixable Far Red Dead Cell Stain (Life Technologies L10119) was used for live/dead discrimination. Data were acquired using Cytek Aurora and analysed in FlowJo (Tree Star). To screen for changes in soluble factors secreted by differentiating organoids in response to cytokines, supernatants were collected at 48 and 96 hours (n=7) after addition of EVTM-NRG1, with or without cytokines. A semiquantitative fluorescent chip-based sandwich ELISA was used to screen for changes in ∼1000 soluble factors in supernatants (human array L-series 1000, slides L-507, L-493 (cat no. AAH-BLG- 1000, L-507 and L-493) RayBiotech). Supernatants were diluted 1 in 7, and assayed according to the manufacturer’s instructions by array testing service (Tebu-bio.com).

#### Single-cell RNA-sequencing of differentiating trophoblast organoids

Single cell suspensions were isolated from six trophoblast lines at selected timepoints following initiation of differentiation in EVTM-NRG1 (Table S3). Specifically, the first three lines (from donors 1, 2, and 3) were collected at 3h, 24h and 48h after the addition of EVTM- NRG1 with and without cytokine treatment. The remaining three lines (from donors 4, 5, 6) were collected before (-48h) and after (0h, 48h and 96h) addition of EVTM-NRG1, with the first two time points (-48h and 0h) collected without cytokine treatment and the other two time points collected from both treatment conditions. For the latter three lines (donors 4, 5, 6), cells grown in TOM supplemented with and without cytokines were also collected at 96h as a control. Multiplexing was performed for each three-donors on the same 10x Genomics reaction.

To next conduct the scRNA-seq experiment, the Chromium Single Cell 3′ Kit v3 from 10X Genomics was used. Single-cell library preparation was performed based on the manufacturer’s protocol to obtain between 2,000 and 10,000 cells per reaction, followed by sequencing on the Illumina HiSeq 4000 or Novaseq 6000 systems to aim for a minimum coverage of 20,000 raw reads per cell.

#### RT-PCR

Trophoblast organoids were plated in Matrigel droplets into 35mm dishes (Ibidi µ-Dish 35 mm #81156) and grown in TOM. When >50% of organoids reached a diameter of 200um, the medium was switched to EVTM. After ∼5 days, when outgrowths of cells were observed from the organoids, the medium was switched to EVTM without NRG1 and cytokines (CSF1, CSF2, CCL5 and XCL1, as detailed above) were added to the medium for a further 96 hours. At different time points during the differentiation process, the organoids and migrating EVT were scraped and collected. For EVT isolation, Matrigel was removed with Cell Recovery Solution (Corning 354253), then the organoids were resuspended in PBS in a low-binding 1.5ml tube and left for 2 minutes for the big organoids to sink to the bottom of the tube. The upper 2/3 of the supernatant, containing EVT detached from the organoids, were collected. This process was repeated three times. Total RNA was extracted using the miRNeasy isolation kit (Qiagen 7004). Total RNA was retro-transcribed using SuperScript^TM^ VILO^TM^ cDNA Synthesis Kit (ThermoFisher, #11754250). The expression of selected transcripts was analysed by quantitative RT-PCR using Taqman gene expression assays (Taqman fast advance Mix, Applied Biosystem, #4444557). Details of the assay IDs for each gene are listed in Table S9. Graphs show expression levels relative to geometric mean of the housekeeping genes *TBP*, *TOP1* and *HPRT1*. The data were processed for statistical analysis with Prism (GraphPad software). Statistical significance calculations were determined using either ratio or paired Student’s *t*-tests. Sample sizes are indicated in the figure legends and were chosen arbitrarily with no inclusion and exclusion criteria. The investigators were not blind to the group allocation during the experiments and data analyses.

#### Immunofluorescence (IF)

Trophoblast organoids were plated in Matrigel droplets into 96 well plates (PhenoPlate Perkin Elmer #6055302) and grown in TOM. When >50% of organoids reached a diameter of 200um, the medium was switched to EVTM. After ∼5 days, when outgrowths of cells were observed from the organoids, the medium was switched to EVTM without NRG1 and cytokines (CSF1, CSF2, CCL5 and XCL1, as detailed above) were added to the medium for a further 96 hours. The organoids were fixed with 4% PFA for 45 min, permeabilized with 0.5% Triton X-100 for 30 min and blocked with 3% donkey serum in PBS for 1h. Primary and secondary antibodies (Table S10) were diluted in PBS with 3% donkey serum and 0.05% Triton X-100 and incubated overnight at 4C. Nuclei were stained with Draq5 (Invitrogen #65-0880-96) together with secondary antibodies incubation. Confocal microscopy was performed on a Leica Stellaris-5 point scanning confocal microscope mounted on a Leica DMI8 inverted microscope equipped with Plan Apochromat 20X/0.75 NA dry objective. Leica LAS X software was used for image acquisitions. For the analysis of MAOA, MCAM and HLA-G intensity, only cells that migrated out of the organoid and whose borders were clearly defined were considered (Figure S7), in order to avoid interference with fluorophores intensity by surrounding or overlapping cells. Cells that met these criteria were individually analysed: ROIs were drawn around individual cells and the specific MAOA, MCAM or HLA-G signals were recorded using ImageJ/Fiji (National Institutes of Health). Integrated density for each cell was corrected for background intensity signal subtracting the integrated density calculated in a cell-free area with the same size. The data were processed for statistical analysis with Prism (GraphPad software). Statistical significance calculations were determined using the non-parametric Mann–Whitney, after assessing the non-normal distribution of the sample with Normal (Gaussian) distribution test. Sample sizes are indicated in the figure legends and were chosen arbitrarily with no inclusion and exclusion criteria. The investigators were not blind to the group allocation during the experiments and data analyses.

#### Immunohistochemistry

The tissue sections from formalin-fixed, paraffin-wax-embedded human implantation sites were dewaxed with Histoclear (National Diagnostics, HS-200), cleared in 100% ethanol and rehydrated through gradients of ethanol (90%, 70%, 50%) to PBS. Epitope retrieval was next conducted in Access Revelation (AR) pH 6.4 (A.Menarini, MP-607-PG1) citrate buffer or Access Super (AS) pH 9 (A.Menarini, MP-606-PG1) Tris-EDTA buffer at 125°C in an Antigen Access pressure cooker (A.Menarini, MP-2008-CE). Sections were then sequentially incubated with 2% blocking serum (of the species where the secondary antibody was made), primary antibodies, biotinylated horse anti-mouse or goat anti-rabbit secondary antibodies, and Vectastain ABC–HRP reagent (Vector, PK-6100). Each incubation lasted for 30 mins at room temperature followed by two washes in PBS. Sections were subsequently developed with diaminobenzidine (DAB) substrate (Sigma, D4168), then counterstained with Carazzi’s haematoxylin, and mounted in glycerol/gelatin mounting medium (Sigma, GG1-10). Images were taken using a Zeiss Axiovert Z1 microscope and Axiovision imaging software SE64 version 4.8. The information of the antibodies used is provided in Table S11.

#### KIR and HLA genotyping

To confirm KIR2DS1+ or KIR2DL1+ status in samples of maternal decidua, genomic DNA was isolated from whole maternal blood samples using the QIAmp DNA mini Blood Kit (Qiagen). From decidual, placental tissue or trophoblast organoids, genomic DNA was isolated following proteinase K digestion as described previously^8^. Genotyping for presence of selected KIR genes or HLA-C was undertaken by PCR-SSP with sequence-specific primers using previously validated methods^18^. C1+ or C2+ HLA-C status in maternal and fetal tissues was determined by high resolution typing of HLA-A/B/C, which was undertaken with an ‘in house’ third generation sequencing pipeline using Pacific Biosciences’ Single Molecule Real-Time DNA sequencing technology as previously described^90^.

#### Analysis of uNK responses to HLA-C

Given the bulk RNA-seq data generated from the three purified uNK subsets after binding to C1+ or C2+HLA-C molecules, we first mapped the sequencing reads to the human reference genome (GRCh38_release37_PRI) using STAR^91^ (version 2.7.8a) by allowing at most one mismatch and requiring the aligned percentage per read greater than 95%. featureCounts from R package Rsubread^92^ was next used to quantify the read abundance for each gene based on the uniquely mapped reads and the human gene annotation (GENCODE v37). We subsequently calculated the reads per kilobase per million mapped reads (RPKM) by normalising the read abundance by total reads mapped and gene length.

To first examine the global effect of KIR binding to C2+ versus C1+HLA-C in each uNK subset, we performed principal component analysis (PCA) using the function “prcomp” in R based on genes with expression level (RPKM) > 1 in all the samples from at least one interaction group (either C1 or C2). To next detect genes differentially expressed (DE) between C2 and C1 groups for each uNK subset, we used the R package “edgeR”^93^ for paired comparisons across groups by the likelihood ratio test based on genes with read counts > 5 in all the samples from at least one group. DE genes were then defined as those with false discovery rate (FDR) < 0.05 and fold change > 1.5. Protein-coding genes exclusively upregulated in KIR2DS1sp uNK after interacting with C2 (no significant upregulation in KIR2DL1sp and KIR2DS1/L1dp uNK) were shown in Figure 1C and Table S12. Functional enrichment analysis was next conducted for those genes using the tool Metascape^94^, with the top 20 enriched terms shown in Figure S1D. The profiles of the common cytokines/chemokines across the decidual cell types as well as their cognate receptors on EVT were based on our previous scRNA-seq data from the first trimester^21^.

#### Preprocessing of scRNA-seq data from organoids

For the scRNA-seq data generated from the organoids treated with and without cytokines, initial preprocessing including read alignment, quality control, and quantification were performed using Cellranger (version 6.0.1) based on the human reference prebuilt in Cellranger (GRCh38-2020-A). Since the organoids derived from different donors were pooled for library preparation and sequencing, Souporcell^95^ (version 2.0) was used to deconvolve the data to derive the donor origin for each cell based on the genotype information of the common variants from the 1000 Genomes. Doublets were further detected by Scrublet^96^ per sample, followed by two rounds of clustering to obtain over-clustered manifold for determining the clusters with significantly higher Scrublet doublet scores based on their median scores^97^ (Benjamini-Hochberg corrected *p*-value < 0.05). Cells that were 1) having > 20% of mitochondria genes; 2) unassigned to a donor or assigned to multiple donors by Souporcell; 3) falling within the doublet-enriched clusters were excluded from the downstream analysis.

#### scRNA-seq data integration, clustering and annotation

Before the data integration, Scanpy^98^ was used to filter genes detected in less than 20 cells, normalise the data (scaling factor 10,000), and detect the highly variable genes (2,000 genes). Data integration across the different donors (six donors) was then performed using scVI^99^, with the “batch_key” set to donors and latent dimension set to 10. The resulting latent representations were then employed for the neighbourhood graph construction, Leiden clustering, and UMAP visualisation. Each Leiden cluster was annotated based on the expression of known marker genes for VCT, SCT and EVT^21,100^ (Figure S3). EVT cells were further subdivided into early EVT (low expression level of HLA-G and remnant expression of VCT genes), intermediate EVT (intermediate expression level of EVT genes), and late EVT (high expression level of EVT genes) subtypes (Figure S3G). For each type of trophoblast cells, there were clusters of cells which were isolated from their corresponding cell types in the UMAP and had a much lower number of genes detected (Figures S3C and S3E). These clusters, together with the cells from the fourth donor which failed to be differentiated to EVT (Figure S3F), were excluded from the following analysis. The data integration was then re-conducted using scVI (1,000 highly variable genes and 10 latent dimensions) after excluding these cells, and the resulting UMAP was shown in Figure 3A.

#### Trajectory analysis in the organoids and primary tissue

scTour^35^ was used to perform all the trajectory analyses. Specifically, a scTour model was first trained based on all the cells from the organoids and 2,000 highly variable genes selected after filtering genes detected in less than 20 cells. The default parameters were used for training. The resulting latent space (alpha_z = 0.7, alpha_predz = 0.3) together with the inferred vector field were used as input for the “plot_vector_field” function to visualise the transcriptomic vector field on the UMAP.

To compare the EVT differentiation pathway between the organoids and the primary tissue, two *in vivo* datasets profiling the maternal-fetal interface using snRNA-seq and scRNA-Seq were considered here^9,21^. A scTour model was first trained based on the EVT cells (VCT_CCC, EVT_1, EVT_2, iEVT, and GC) from the first dataset (snRNA-seq). The intersection of genes expressed in this dataset and organoids with the 500 highly variable genes from the second *in vivo* dataset (scRNA-seq)^21^ were used for model training (473 genes). The resulting model was then employed to infer the developmental pseudotime of the EVT cells from the first dataset, as well as to predict the pseudotime for the *in vitro* EVT cells in the organoids and for the *in vivo* EVTs collected from placenta and decidua in the second dataset. The gene expression dynamics along the pseudotime was then visualised using heatmap.

#### Differential abundance analysis in the organoids

To test the cell abundance changes induced by the cytokine treatment, the R package miloR was used^101^. In detail, the latent space from the scVI model as described above was used to build the neighbourhood graph, with the k-nearest neighbours set to 15. The differential abundance testing was performed between treatment conditions, with samples collected from different donors and time points considered as replicates. Each neighbourhood was annotated as the cell type most dominant within it, that is, more than 70% of the cells in this neighbourhood were from this cell type. Neighbourhoods with significant abundance changes were defined as those with Spatial FDR (multiple testing corrected *p*-value) < 0.1.

#### *In vivo*-*in vitro* scRNA/snRNA-seq data alignment

Three different approaches were adopted to map the cell types from the organoids (scRNA-seq) to those from the two *in vivo* datasets described above (snRNA-seq and scRNA-seq)^9,21^. The first approach used CellTypist^102^ to train a logistic regression classifier based on the *in vivo* trophoblast cells from the first dataset (snRNA-seq). The resulting model was subsequently applied to predict the identities of the cells from the organoids. The second approach assigned each cell from the organoids to the most similar *in vivo* cell type from the second dataset (scRNA-seq) based on their transcriptome similarity. Specifically, the *in vivo* EVT cells collected from the placenta and decidua were firstly separated and labelled as two distinct subtypes. For each gene, the average expression across each *in vivo* trophoblast cell subtype was then computed. This was followed by the calculation of the Pearson correlation coefficient between each *in vitro* cell and each *in vivo* cell subtype based on the highly variable genes. The identities of the *in vitro* cells were determined as the *in vivo* cell subtype with the highest coefficient. The third approach used scVI to integrate the EVT cells from the organoids and the second *in vivo* dataset. For the *in vivo* EVT cells, the four donors D9, D10, D11, D12 which had more than 100 EVT cells collected were included here for the integration. The model training was based on 1,000 highly variable genes, with the “batch_key” set to donors and latent dimensions set to 10. The latent space derived from the model was then used to yield the Leiden clusters and UMAP visualisations for the *in vitro* and *in vivo* EVT cells as shown in Figure S4B.

#### Differential expression analysis in the organoids and primary tissue

To detect gene expression changes induced by uNK cytokines in the organoids, the Wilcoxon test followed by Bonferroni correction were applied through the function “rank_genes_groups” within Scanpy for each cell subtype. Genes with corrected *p*-value less than 0.05 were considered to be statistically significant. Here we focused on the molecular changes in EVT_late_3 subtype which highly resembled the *in vivo* iEVT and was most affected by cytokines. To further examine their profiles *in vivo* between EVT located at the proximal end of the cell column and iEVT, we considered the two *in vivo* datasets described above (snRNA-seq and scRNA-seq)^9,21^. Specifically, Wilcoxon test was performed between EVT cells located in the cell column (VCT_CCC and EVT_1) and those in deeper decidua (iEVT and GC) (snRNA-seq), as well as between EVT cells collected from placenta and those from decidua (scRNA-seq). The final genes with expression affected by cytokines in the organoids meanwhile showing *in vitro*-*in vivo* consistent changes were defined using three criteria: 1) corrected *p*-value calculated in EVT_late_3 subtype between cells treated with and without cytokines < 0.05; 2) corrected *p*-value estimated in the first *in vivo* dataset between EVT cells from cell columns and deep decidua < 0.05; 3) corrected *p*-value computed in the second *in vivo* dataset between placental and decidual EVT cells < 0.05; 4) consistent direction of changes across the three comparisons. The functional enrichment analysis for the genes identified was performed by Metascape^94^.

#### Analysis of cellular interactions between EVTs and surrounding populations

To assess the potential effects of cytokines on the downstream cellular interactions, we used CellphoneDB^103,104^ (version 3.1.0) to identify the ligand-receptor based communication between decidual EVTs and their surrounding populations including uNKs, macrophages, perivascular cells, stromal cells, and endothelial cells. Specifically, the expression matrix and meta information for these cell types from the *in vivo* scRNA-seq dataset^21^ were used as input for CellphoneDB to perform the statistical analysis. The significant ligand-receptor pairs mediating the interaction between EVTs and a certain cell population were defined as those with *p*-value less than 0.05. The pairs focusing on the cytokine receptors that were affected by the cytokine treatment were shown in Figure S5E.

## Supplemental Figures

**Figure S1.**
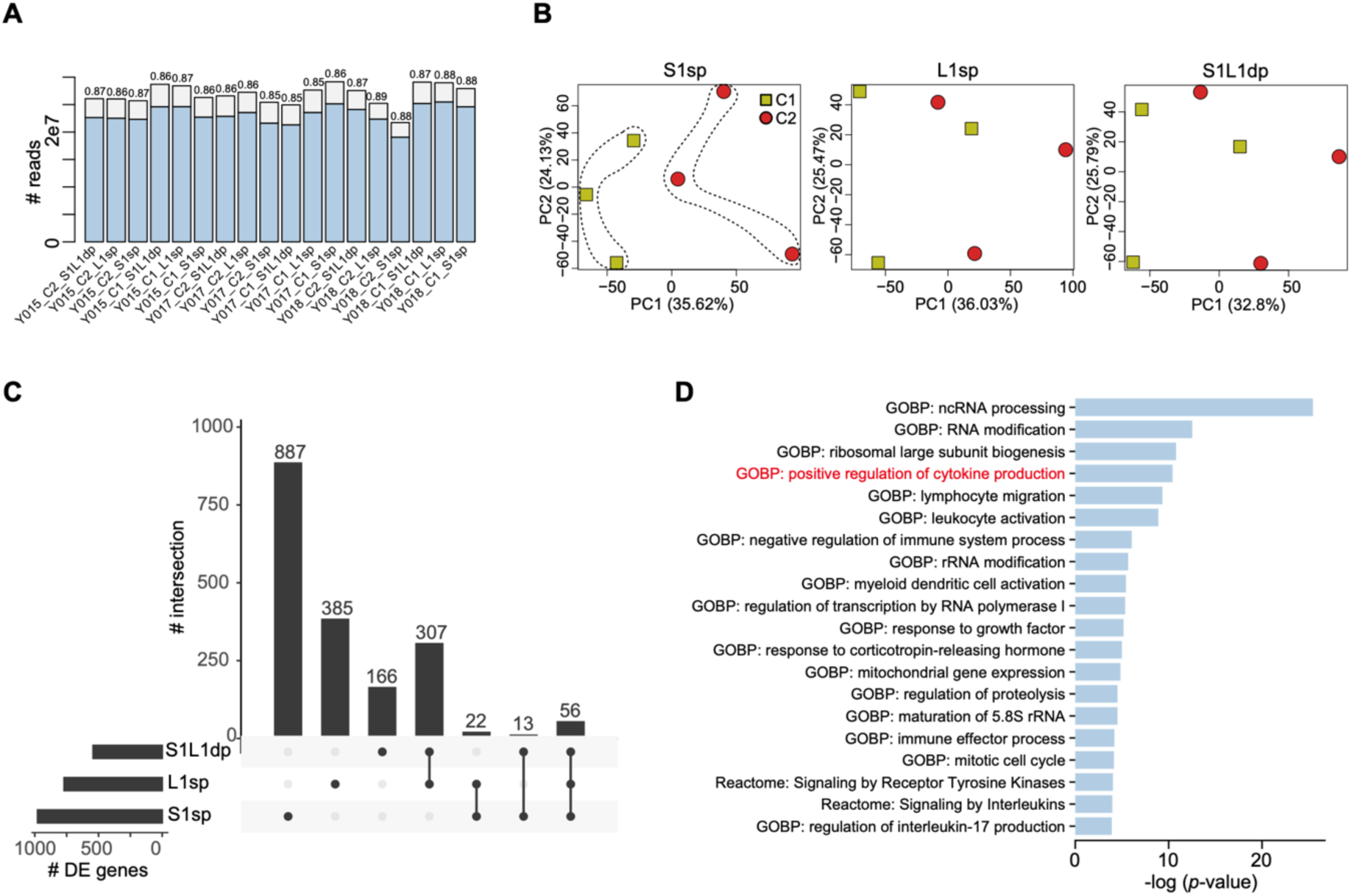
Responses of KIR+ uNK subsets to HLA-C. (**A**) The number of total reads obtained from each sample, with the number of reads uniquely mapped to the reference genome shown in blue and the corresponding percentage on top of each bar. (**B**) The first two principal components (PCs) of principal component analysis (PCA) for each uNK subset. Dots indicate samples, with colours representing binding to 221-C2+ or 221-C1+ targets. (**C**) The number of genes differentially expressed (DE) between C2 and C1 of HLA-C in each uNK subset (FDR < 0.05 and fold change > 1.5, left), with DE genes shared with the other subsets shown in top bars. (**D**) GO biological processes and Reactome pathways enriched in genes showing specific upregulation in KIR2DS1+ uNK.

**Figure S2.**
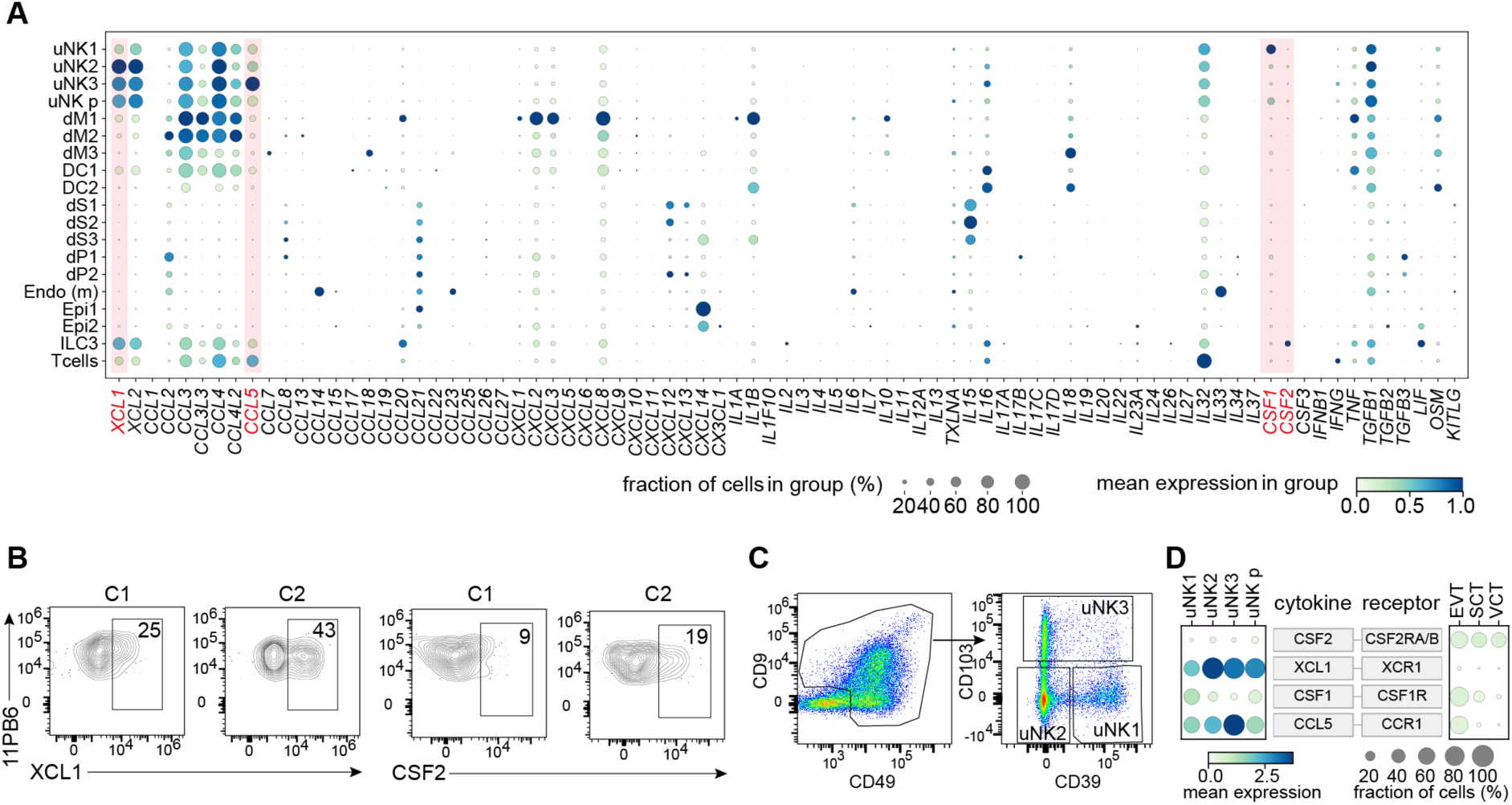
Expression of cytokines/chemokines in scRNA-seq data from maternal-fetal interface. (**A**) Dot plot displaying the expression of common cytokines/chemokines in decidual cell populations based on our previous scRNA-seq data. The colour gradient denotes the maximum-normalised expression level and dot size represents the proportion of cells expressing the genes. dM, decidual macrophages; dS, decidual stromal cells; Endo, endothelial cells; Epi, epithelial glandular cells; dP, perivascular cells; DC, dendritic cells. (**B**) Intracellular flow cytometry confirming upregulation of XCL1 and CSF2 in KIR2DS1sp uNK cells after co-culture with 221-C2+HLA-C compared with 221-C1+HLA-C targets (representative sample from n=6 individuals). Gating of KIR2DS1sp uNK cells is shown in Figure 1B. (**C**) Gating strategy for uNK1-3 subsets. (**D**) Dot plot displaying the expression of the four cytokines (CSF2, XCL1, CSF1, and CCL5) in uNK subsets and their corresponding receptors in decidual EVT based on our previous scRNA-seq data. The colour gradient denotes the log-transformed normalised expression level and dot size represents the proportion of cells expressing the genes. p, proliferative.

**Figure S3.**
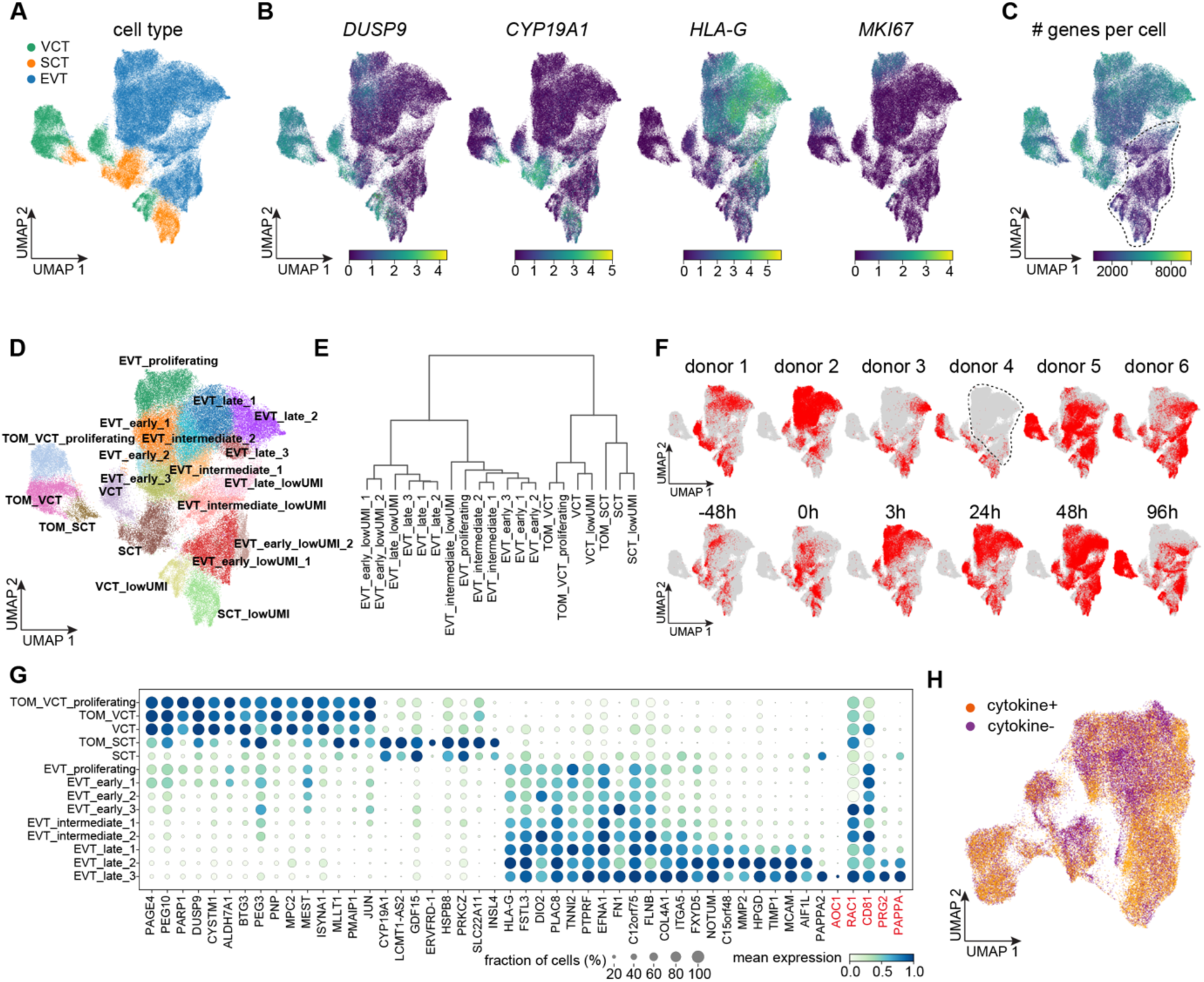
Quality control of the single-cell data from trophoblast organoids. (**A**) UMAP visualisations of the 94,752 cells from trophoblast organoids with colours indicating cell types. (**B**) UMAP visualisations of expression of marker genes for each cell type (*DUSP9*, *CYP19A1*, *HLA-G*) and proliferation (*MKI67*). (**C**) UMAP visualisations of the number of genes detected in each cell. The dotted circle indicates cells with a low number of genes detected. (**D**) UMAP visualisations of the cell subtypes. (**E**) Unsupervised hierarchical clustering of the cell subtypes from (**D**). (**F**) UMAP visualisations of the cells from different donors and time points. The dotted circle indicates the absence of EVT in donor 4. (**G**) Dot plot displaying the expression of established trophoblast marker genes across cell subtypes identified from the organoids. Genes highlighted in red represent GC marker genes. The colour gradient denotes the maximum-normalised expression level and dot size represents the proportion of cells expressing the genes. (**H**) UMAP visualisations of the 67,996 high quality cells coloured by treatment conditions.

**Figure S4.**
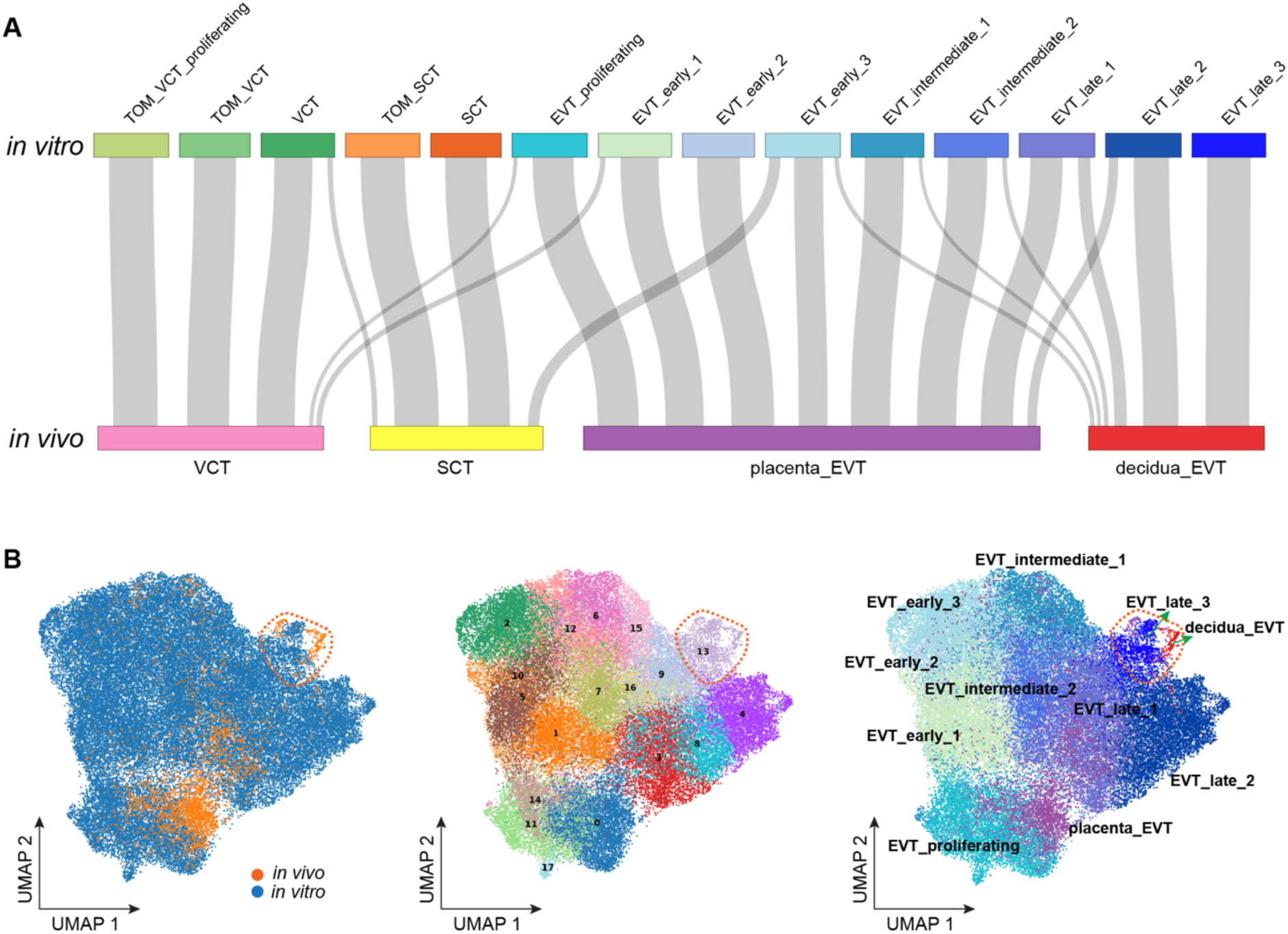
Alignment of *in vitro*-*in vivo* trophoblast cell types. **(A)** Sankey plot showing the correspondence between the cell subtypes identified from the organoids (top) and those from the primary tissue (bottom). (**B**) Integration of *in vitro* and *in vivo* EVT cells, with cells colored by biological system (left), Leiden clusters (middle), and cell annotations (right). The dotted circle highlights the cluster 13 formed by *in vitro* EVT_late_3 subtype and *in vivo* EVT cells collected from decidua.

**Figure S5.**
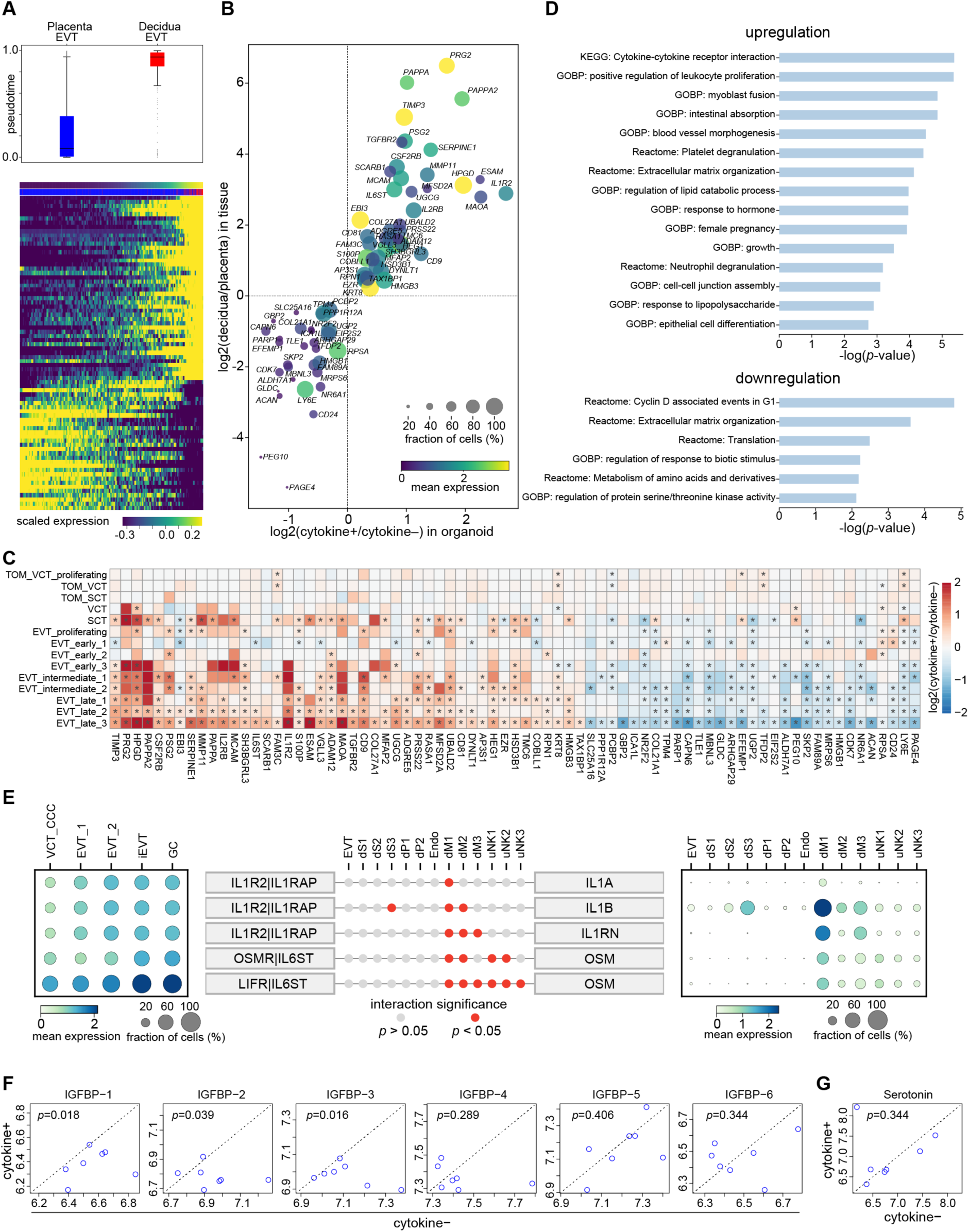
The diverse effects of uNK cytokines on EVT. **(A)** Top panel: box plot of the developmental pseudotime inferred for the *in vivo* EVT cells collected from placenta and decidua based on the same scTour model as in Figure 3E. The medians, interquantile ranges, and 5th, 95th percentiles are indicated by centre lines, hinges, and whiskers, respectively. Bottom panel: the expression dynamics along the developmental pseudotime for genes shown in Figure 3F. (**B**) Scatter plot showing genes consistently upregulated or downregulated after cytokine treatment in EVT_late_3 in organoids (x axis) and after invading the decidua (decidual EVT) compared to EVT collected from placenta based on the scRNA-seq dataset (y axis). Colour shade and dot size denote the expression and proportion averaged between the *in vitro* and *in vivo* EVT, respectively. (**C**) Heatmap showing the log2-transformed fold change between cells treated with and without cytokines in each cell type for genes shown in (**B**). Asterisks indicate significant changes (Bonferroni corrected *p*-value < 0.05). (**D**) GO biological processes, KEGG and Reactome pathways enriched in upregulated and downregulated genes shown in (**B**). (**E**) Ligand-receptor interactions between decidual EVT cells and surrounding populations. Statistically significant interactions (*p*-value < 0.05) are highlighted in red. The expression patterns of the receptors across the EVT subtypes, and the ligands across the decidual populations are shown in the left and right panels, respectively. The colour gradient denotes the log-transformed normalised expression level and dot size represents the proportion of cells expressing the genes. (**F**) Log-transformed relative levels of IGFBP in supernatant of organoids treated with cytokines (y-axis) and without cytokines (x-axis), measured using raybiotech cytokine arrays. The *p*-value calculated between the two conditions by one-sided Wilcoxon test is shown on top. (**G**) As with (**F**), but for serotonin.

**Figure S6.**
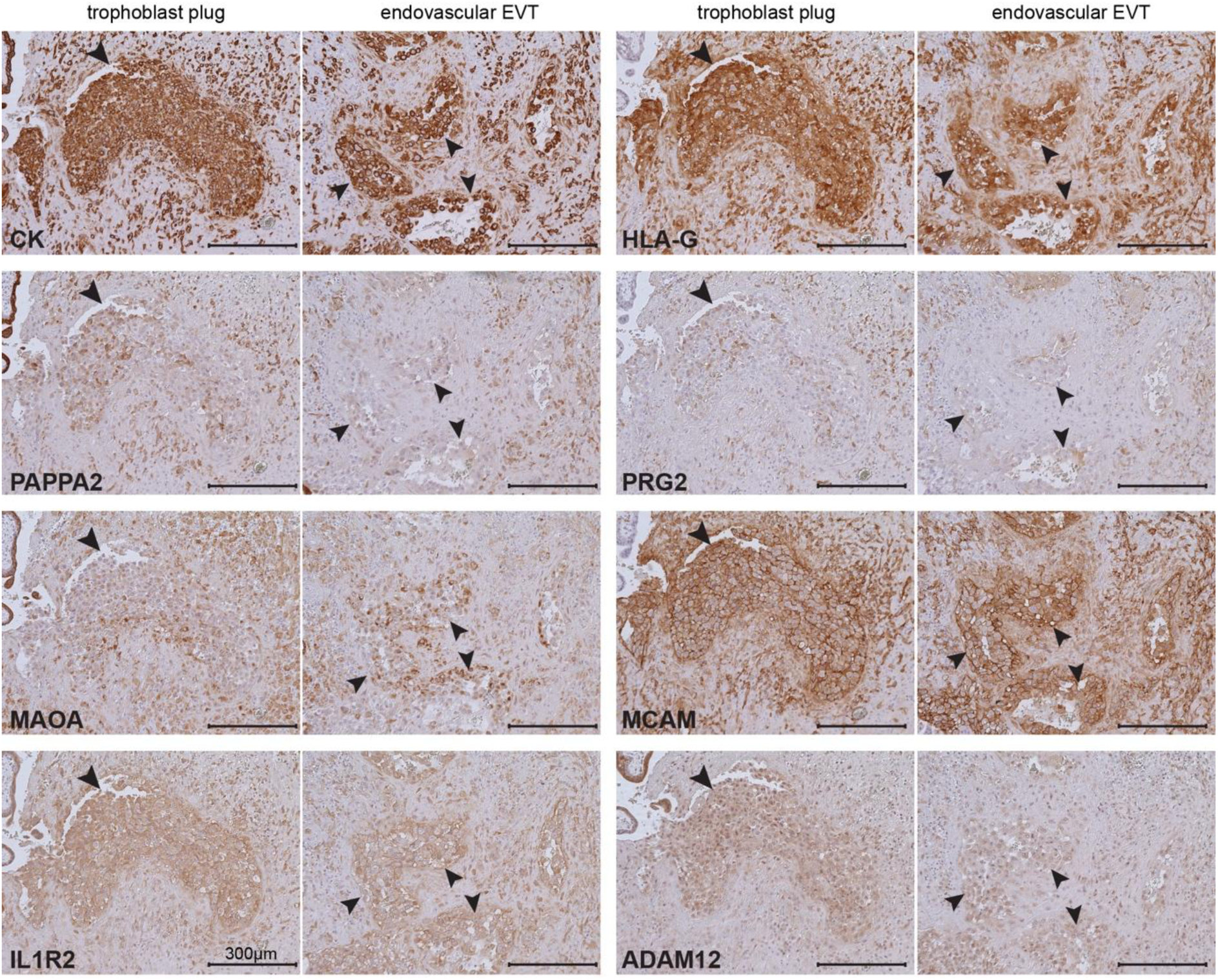
Expression pattern of uNK cytokine-regulated genes in endovascular EVT. Immunohistochemistry of sections of endovascular EVT in trophoblast plugs (left) and within arteries (right) for proteins highlighted in red in Figure 4A together with Cytokeratin (CK) and HLA-G. Arrows indicate the trophoblast plug (left) and endovascular EVT (right).

**Figure S7.**
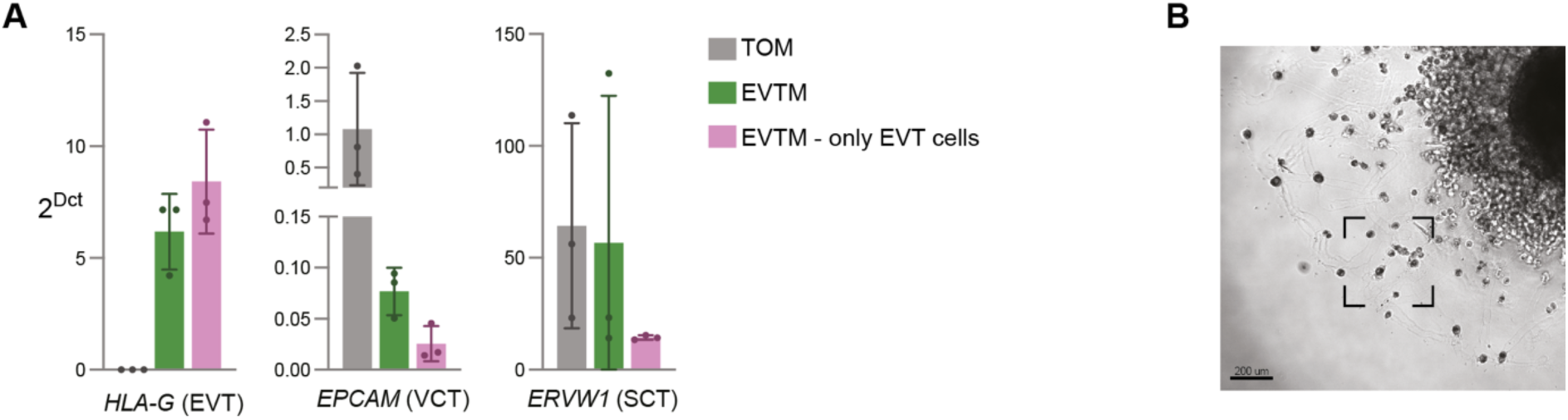
Isolation of EVT cells for gene and protein expression analysis. **(A)** RT-PCR of trophoblast sub-population markers performed in trophoblast organoids, comparing RNA extracted from organoids maintained in TOM, organoids differentiated with EVTM, or EVT cells isolated from organoids differentiated with EVTM. Results are expressed as 2^-Δct^. Bars represent mean ± SD (*n*=3 independent experiments). (**B**) Representative brightfield image of EVTM-differentiated trophoblast organoid. The square indicates the EVT cells migrating out from the organoid, representing the cells analysed by IF in Figure 5C.

**Table S1. Genes associated with disorders of pregnancy.**

**Table S2. Composition of Trophoblast Organoid Medium (TOM).**

**Table S3. Metadata of trophoblast organoids and decidual cell donors.**

**Table S4. Antibody panel for uNK cell phenotyping.**

**Table S5. Antibodies for cell sorting of KIR2DS1+ and KIR2DL1+ uNK subsets.**

**Table S6. Antibodies for intracellular cytokine staining to monitor responses in uNK subsets.**

**Table S7. Composition of medium for the differentiation of EVT (EVTM) from trophoblast organoids.**

**Table S8. Antibodies for cytokine receptor staining in primary EVT or trophoblast organoids.**

**Table S9. Genes analysed by RT-PCR.**

**Table S10. Antibodies used in immunofluorescence.**

**Table S11. Antibodies used in immunohistochemistry.**

**Table S12. Protein-coding genes specifically upregulated in KIR2DS1sp uNK after binding to C2+HLA-C.**

## Notes

### Competing Interest Statement

The authors have declared no competing interest.

